# Conformational dynamics in the membrane interactions of bispecific targeted degrader therapeutics

**DOI:** 10.1101/2025.03.18.643966

**Authors:** Emma Inganäs, Hanna Lavén, Rémi Caraballo, Fredrik Klingegård, MyLan Eklund, Linda Andersson, Constanze Hilgendorf, Pär Matsson

## Abstract

Proteolysis targeting chimeras (PROTACs) offer vast new therapeutic opportunities, however their physicochemical properties are difficult to combine with optimal cell permeability and exposure at the target sites. We have systematically analyzed a dataset of more than 3500 PROTACs to investigate how the choice of ubiquitin E3 ligase ligands, linker design, and global molecular properties can be optimized to achieve the desired cell permeability and intracellular exposure. We find that conformational flexibility leads to environment-dependent shielding of polar functions and improved interactions with cell membranes, but that at the same time extended, linear conformations within the membrane are beneficial. Linker composition was a major factor in determining the folding propensity. Collectively, our results suggest that strategies to rationally design linkers and to shield polarity selectively within the protein-of-interest (POI) ligand and/or E3 ligand domains, rather than more extensive folding, may be beneficial in the design of permeable and effective PROTACs.

## Introduction

Targeted protein degradation has become a valuable new tool in the drug discovery and chemical biology toolbox. Targeted degraders, or proteolysis targeting chimeras (PROTACs), work by hijacking the cell’s ubiquitin-proteasomal degradation system: bispecific ligands simultaneously bind the target protein and an E3 ligase, and the induced proximity leads to selective ubiquitination and subsequent degradation of the target. ^1–8^

PROTACs have unique characteristics that set them apart from traditional therapeutic modalities. ^9^ Most importantly, they circumvent the need for constant target occupancy: conventional drugs are only effective when all or most target molecules are occupied, and when these are bound in a way that alters the function of the target protein. In contrast, a single PROTAC molecule can iteratively bind to, and induce degradation of multiple copies of the target protein in a substoichiometric, catalytic fashion. ^2,10^

As with all drugs that target intracellular biomolecules, PROTACs must be able to cross the plasma membrane to elicit the therapeutic response. Importantly, not all of the drug that enters the cell interior is available to bind to intracellular targets. Typically, a variable and unpredictable fraction of the drug is bound to cellular components or trapped in cellular organelles. ^11–13^ We have previously shown that *free* intracellular drug concentrations are an excellent predictor of intracellular drug potency for various intracellular targets, ^11–13^ and that the pharmacologically active free intracellular drug fraction is critically determined by non-specific binding to cellular phospholipid membranes. ^11,14,15^

The bispecific affinity of the degrader molecule leads to a considerably greater molecular size than in traditional, cell permeable molecules, significantly restricting intracellular exposure. ^16^ Furthermore, the larger size of PROTACs is typically accompanied by corresponding increases in additional physicochemical descriptors, such as lipophilicity and the number of hydrogen bonding functionalities, ^16^ often considerably overstepping the boundaries of traditional, rule-of-5 compliant drug-like chemical space. Thus, PROTACs are situated in a challenging region of chemical space, where large size in combination with polar functionalities may limit cell permeability, whereas more lipophilic PROTACs can exhibit both problematic solubility, and excessive binding to cellular and subcellular lipid membranes, potentially affecting their pharmacokinetics and pharmacological activity. ^17^

We previously demonstrated that cell permeability in macrocyclic molecules^18^ and in orally available beyond-rule-of-5 drugs^19,20^ is improved by, and sometimes contingent on, the ability to adaptively expose or shield molecular features depending on the surrounding environment. Such environment-dependent conformational sampling has since been demonstrated in specific case studies also for PROTACs, indicating that similar rules govern their cell permeability as for other beyond-rule-of-5 molecules. ^21,22^

Numerous studies have demonstrated that the length and type of linker connecting the two protein ligands can have dramatic effects on degradation efficiency. ^1,23,24^ In addition, the choice of linker chemistry and length has been shown to lead to large differences in permeability in structurally similar PROTAC series, ^25,26^ and that permeability differences may be associated with conformational flexibility in the linker region. ^21,22^ However, whether these initial findings generalize to other PROTACs and can be used in prospective drug design remains unknown.

We have, therefore, collected and analyzed a considerable proportion of the PROTACs available to date in the public domain, with the aim to define structural features and molecular properties that govern their interactions with, and transport across cellular membranes.

## Results

### Physicochemical characteristics of PROTACs

A database comprising 3576 PROTAC molecules was collated from public data sources, and complemented with recent PROTACs literature. ^16,27^ Of these, 2304 (64%) contained a cereblon-binding domain (CRBN), 1195 (33%) contained a Von Hippel-Lindau-binding domain (VHL) (**Table 1**) and the rest were reported to bind to other E3 ligases (**Supplementary Table 1**). Additionally, 2.4% of the VHL-binding PROTACs had CRBN as the degradation target and thus also contained a CRBN-binding POI ligand.

**Table 1.** Molecular properties of literature PROTACs. Summary of molecular descriptors (median (min-max range)) of full PROTAC molecules, and for the separated E3 ligand, linker and POI ligand domains for CRBN-binding and VHL-binding PROTACs.

| CRBN-binding PROTACs |  |  |  |  |
| --- | --- | --- | --- | --- |
|  | Full molecule | E3 ligand | Linker | POI ligand |
| **n types** | 2304 | 60 | 1174 | 444 |
| **MW<sup>a</sup> [Da]** | 874 (409-2009) | 272 (188-511) | 182 (0-647) | 423 (60-1581) |
| **cLogP<sup>b</sup>** | 4.4 (-7.9-12.3) | 0 (-1.3-3.4) | 0.7 (-1.7-7) | 3.8 (-6.9-8.6) |
| **TPSA<sup>c</sup> [Å<sup>2</sup>]** | 214 (87-793) | 96 (46-175) | 37 (0-150) | 82 (3-654) |
| **HBD<sup>d</sup>** | 4 (1-25) | 2 (0-2) | 1 (0-3) | 2 (0-23) |
| **HBA<sup>e</sup>** | 13 (4-30) | 5 (2-11) | 3 (0-12) | 6 (1-24) |
| **nRotB<sup>f</sup>** | 17 (4-62) | 2 (1-8) | 8 (0-38) | 5 (0-52) |
| VHL-binding PROTACs |  |  |  |  |
|  | Full molecule | E3 ligand | Linker | POI ligand |
| **n types** | 1195 | 74 | 543 | 256 |
| **MW<sup>a</sup> [Da]** | 1051 (717-1684) | 458 (390-705) | 162 (0-777) | 418 (167-989) |
| **cLogP<sup>b</sup>** | 7.1 (0.9-14.4) | 2.4 (0.4-4.6) | 1 (-1.3-5.6) | 4 (-2.4-9) |
| **TPSA<sup>c</sup> [Å<sup>2</sup>]** | 217 (124-356) | 112 (91-152) | 24 (0-149) | 80 (12-244) |
| **HBD<sup>d</sup>** | 5 (2-10) | 3 (2-4) | 0 (0-2) | 2 (0-7) |
| **HBA<sup>e</sup>** | 14 (8-25) | 6 (4-11) | 2 (0-14) | 6 (1-12) |
| **nRotB<sup>f</sup>** | 21 (5-44) | 6 (5-12) | 8 (0-32) | 5 (0-18) |

<sup>a</sup> molecular weight; <sup>b</sup> calculated LogP; <sup>c</sup> topological surface area; <sup>d</sup> hydrogen bond donors;
<sup>e</sup> hydrogen bond acceptors; <sup>f</sup> number of rotatable bonds

VHL-binding PROTACs were, on average, larger and more lipophilic (as assessed by their molecular weight, MW, and octanol-water partition coefficients, cLogP) (**Table 1**). The number of hydrogen bond acceptors and donors (HBA/HBD) were similar in VHL- and CRBN-binding PROTACs and thus it follows that their topological polar surface areas (TPSA) were approximately equal. Consequently, the increased size results mostly from non-polar atoms, explaining the substantially greater lipophilicity in VHL-binding PROTACs (**Table 1**).

By separating each of the compiled PROTAC structures into their three functional parts (**Fig. 1B**), we could assess the relative contributions from each to the properties of the whole molecule. Excluding stereochemical variations, a total of 127 different E3 ligands, 1540 different linkers and 591 different POI ligands were identified in the dataset. While POI ligands and linkers were similar in size in all classes of PROTACs, the greater overall size of VHL-directed compared to CRBN-binding PROTACs originated primarily from their substantially larger E3 ligands.

**Figure 1.**
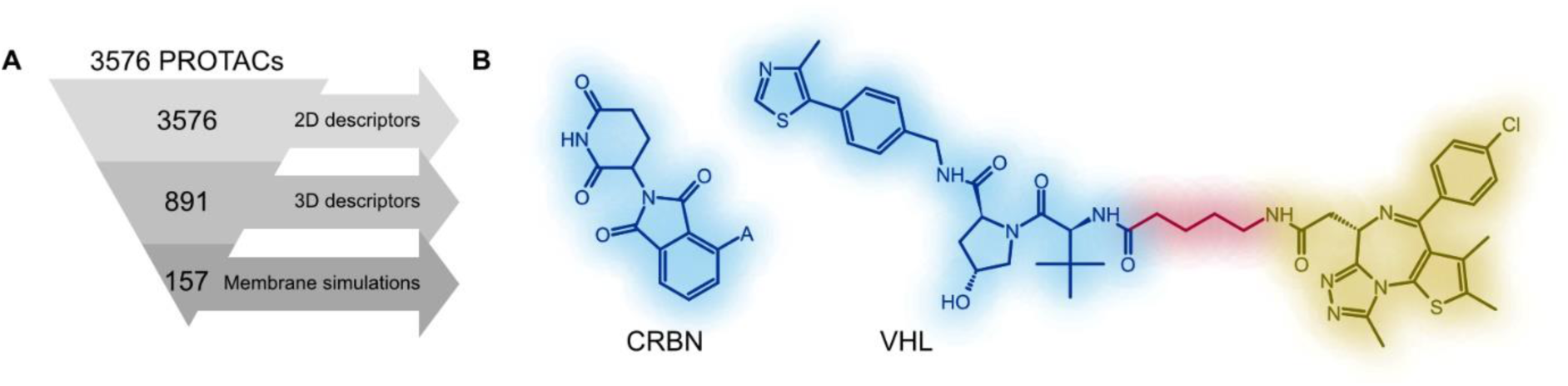
A) Overview of number of PROTAC molecules included in each stage of the analysis. B) Molecular structures for prototypical CRBN- and VHL-binding PROTACs.

On average, the POI ligand was the most lipophilic part of the PROTAC, and VHL-binding domains more lipophilic than CRBN-binding domains. The VHL-binding domains also contained more rotatable bonds, indicating greater conformational flexibility than in CRBN ligands. Notably, HBA, HBD and TPSA were similar for the different E3 ligand domains, despite their considerable size difference. These polarity descriptors were also similar for the POI ligand domains in VHL- and CRBN-binding PROTACs.

### Conformation Dependent Polarity Shielding

We previously demonstrated that large and conformationally flexible (‘beyond-rule-of-5’) molecules dynamically expose or shield polar atoms depending on the surrounding environment. ^18,20^. This has since also been demonstrated in smaller case studies of PROTACs. ^21,22,28,29^ To assess if such conformation-dependent polarity is a recurrent feature in PROTACs and how it in turn affects membrane affinity and permeation, we systematically analyzed almost 900 chemically diverse PROTACs representative of the full PROTAC collection. The subset was subjected to extensive molecular mechanics conformational sampling in water and chloroform environments to explore the structural dynamics of PROTACs in relevant biological compartments.

Fewer distinct low-energy conformations were identified in chloroform than in water for the vast majority of all PROTACs, indicating a restricted conformational sampling in the non-polar, membrane-mimicking solvent (**Supplementary Table 2**). Since these were implicit-solvent calculations, the different conformational diversity reflects constraints arising from the electronegativity of the respective solvents rather than steric effects and lateral pressure that would also contribute within a real lipid membrane environment. Our results were consistent with NMR-derived conformations for a PROTAC molecule reported by Atilaw et al., ^21^ for which a limited number of conformations were observed in each of the three studied solvents (chloroform, DMSO, and DMSO-D_2_O), with greater conformational variations in the polar solvents, and with little overlap between the respective conformation ensembles.

In general, the conformations exposed greater solvent-accessible PSA in water than in chloroform, irrespective of which E3 ligand domain the PROTAC contained. Further, CRBN-binding PROTACs exposed more polar surface in either solvent than did VHL-binding ones (**Supplementary Table 2**), despite containing similar numbers of polar atoms and having similar topological PSA. It thus appears that VHL-binding PROTACs are able to hide more polar surface area than CRBN-binding ones.

To further understand the degree of conformation-dependent polarity shielding in PROTACs, the difference between the maximum and the minimum PSA among each molecule’s conformations was calculated, and normalized to the maximum PSA (to account for between-compound differences in the ‘shieldable surface area’ due to their different polar atom content). The within-compound ranges of the solvent-exposed PSA varied between 2% and 80% of the maximum exposed PSA for CRBN-binding PROTACs, and up to 300 Å^2^ (6-300 Å^2^) PSA was found to be hidden in the most shielded conformations. For VHL-binding PROTACs these numbers were instead between 14% and 76%, with up to 248 Å^2^ (25-248 Å^2^) PSA shielded. Thus, we conclude that polarity shielding is a common feature of PROTACs regardless of which E3-ligase is targeted, but that the ability to shield polar groups varies greatly (**Fig. 2A-D**).

**Figure 2.**
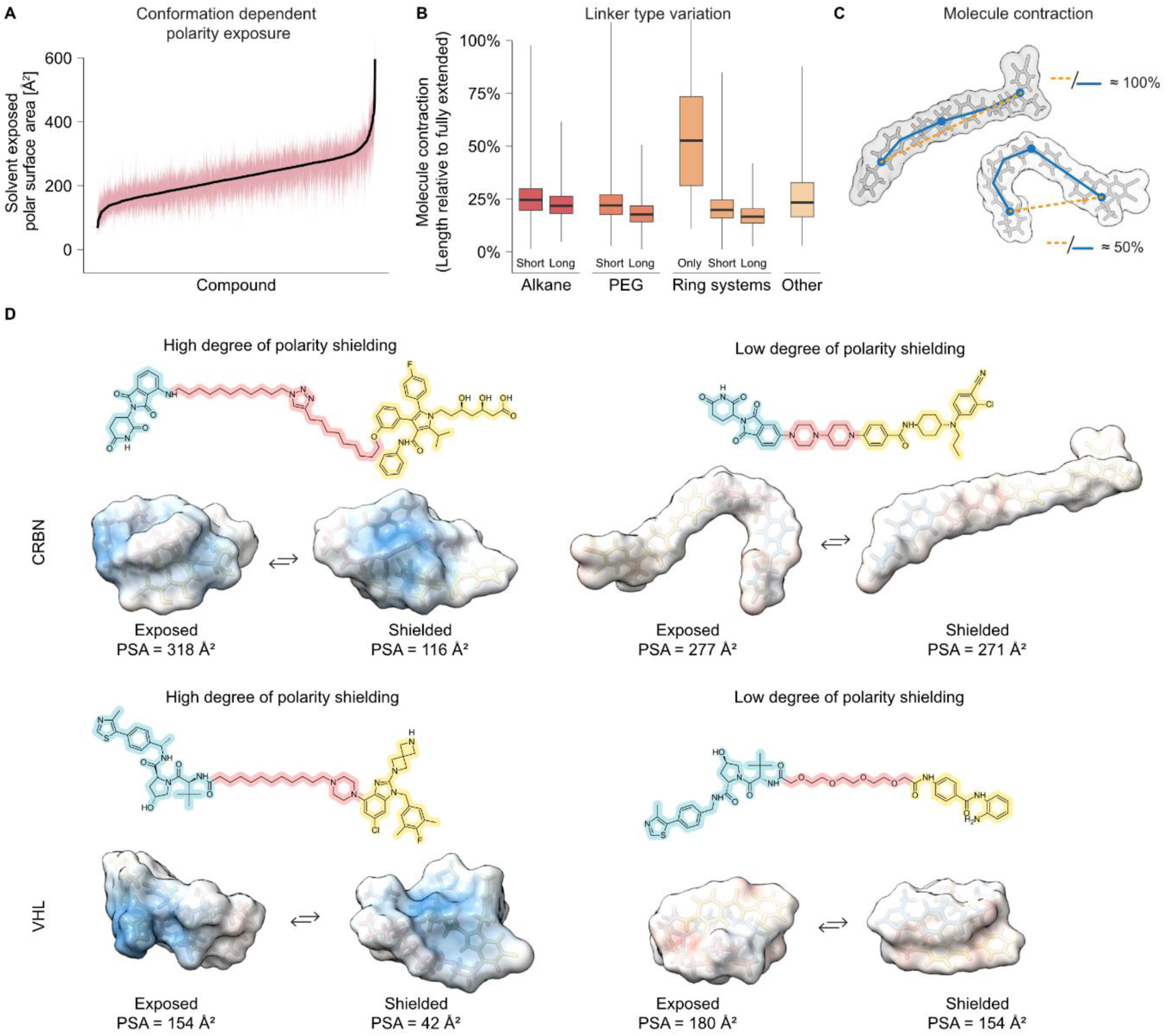
Environment-dependent exposure of polar atoms and molecule contraction. A) Large variability is seen in the ability of different PROTACs to shield polar surface area. Black solid line indicates median and red shaded area indicates min-max range (n = 891). B) Differences in molecule contraction among the different linker type groups (n = 891 PROTACs). Black solid line indicates the median and the bottom and upper box limits are the 25^th^ and 75^th^ percentile. Whiskers indicate the minimum and maximum values. C) Explanation of the molecule contraction parameter. D) Examples of CRBN-binding (top panels) and VHL-binding (bottom panels) molecules with high degree (left panels) and low degree (right panels) of shielded polar surface area.

### Linker characteristics and conformational flexibility

While the POI ligand and E3 ligand domains have clear and direct roles in the proximity-inducing effect, it is becoming clear that also the linker region can markedly affect degradation efficiency by steering the distance and relative orientation in the ternary complex. ^1,23,24^ Much less is known of how linker chemistry and structure affect membrane interactions and cellular exposure.

To enable delineation of such impact, the separated linker structures were further analyzed to identify commonly occurring substructures. Visual inspection yielded a list of motives, including saturated alkyl and polyethylene glycol (PEG) chains of increasing lengths, discrete substructures such as azo, alkyne and sulfur-containing moieties, and various ring systems, which were subsequently mapped onto each of the identified linkers using a custom RDKit script. The most common linker substructures were alkyl chains (found in approximately 37% of all linkers) and PEG chains (found in one third of all linkers) (**Supplementary Fig. 1**).

Frequencies of both linker motifs decreased with increasing motif length. Further, 47% of the linkers contained at least one amide bond and 56% contained some kind of ring system, the most common of which were piperazines and triazoles (**Supplementary Fig. 1**). Notably, the predominance of simple aliphatic and PEG-based linkers in reported PROTACs likely results from a bias in the literature towards early-stage discovery projects primarily aimed at identifying suitable POI ligands and connection vectors, whereas reported clinical-stage PROTACs often contain shorter, more rigid ring-containing linkers. In our collated literature dataset such linkers were less frequent and observed in 6% of the molecules.

Conformational flexibility in PROTACs can arise from rearrangements in any combination of the three constituent parts—the E3 ligand, linker and POI ligand domain—as well as in their movement relative to each other. To assess the underlying dynamics leading to variable conformational sampling in different PROTACs, we calculated two complementary parameters: ‘linker contraction’, which describes the fractional reduction in linker length compared to its fully extended form, and ‘molecule contraction’, which describes how the entire structure folds by comparing the distance between the centers-of-mass for the E3 ligand and POI ligand domains in relation to the fully extended structure (**Supplementary Fig. 2**).

We characterized the ability of each of the different linker classes to form folded structures and found that PEG-based linkers of length ≥ 3 monomers (i.e., ≥ 9-atom chains), with or without ring systems, gave the most folded molecules (**Fig. 2B**). This was consistent with NMR-derived conformations for two PEG-linked PROTACs, where longer chains were shown to favor the gauche effect and a more folded structure ^22^. In contrast, linkers consisting solely of rings yielded conformations that, on average, were close to fully stretched (**Fig. 2B**). Alkyl-based linkers and ring systems connected through short linear chains resulted in modest folding.

Notably, reduction in exposed PSA, correlated stronger with an overall folding of the molecules (R^2^ = 0.02-0.26, **Supplementary Fig. 1**) than with contraction of the linker region (R^2^ = 0-0.06), indicating that polarity shielding results from conformational changes that extend outside the linker itself, and that also involves rearrangement in the POI ligand and ligase domains. Thus, the impact of changing linker chemistry on the overall molecule properties will likely depend strongly on the chemical nature of the E3 ligand and POI ligand domains.

### Interaction with model cell membranes

Permeation across cell membranes is essential for PROTACs to reach their sites-of-action, and binding to subcellular membranes also governs how much of the internalized PROTAC is freely diffusing in the cytosol and available to form the active ternary complex. To examine the relationship between PROTAC chemistry and their interactions with cell membranes more closely we sourced a chemically diverse, representative subset (**Fig. 3A**) from commercial vendors and measured their binding to HEK293 cell homogenates using equilibrium dialysis. ^11,12^ Matched molecular pairs were included in which POI and E3 ligands were kept constant, and the length or chemistry of the linker varied.

**Figure 3.**
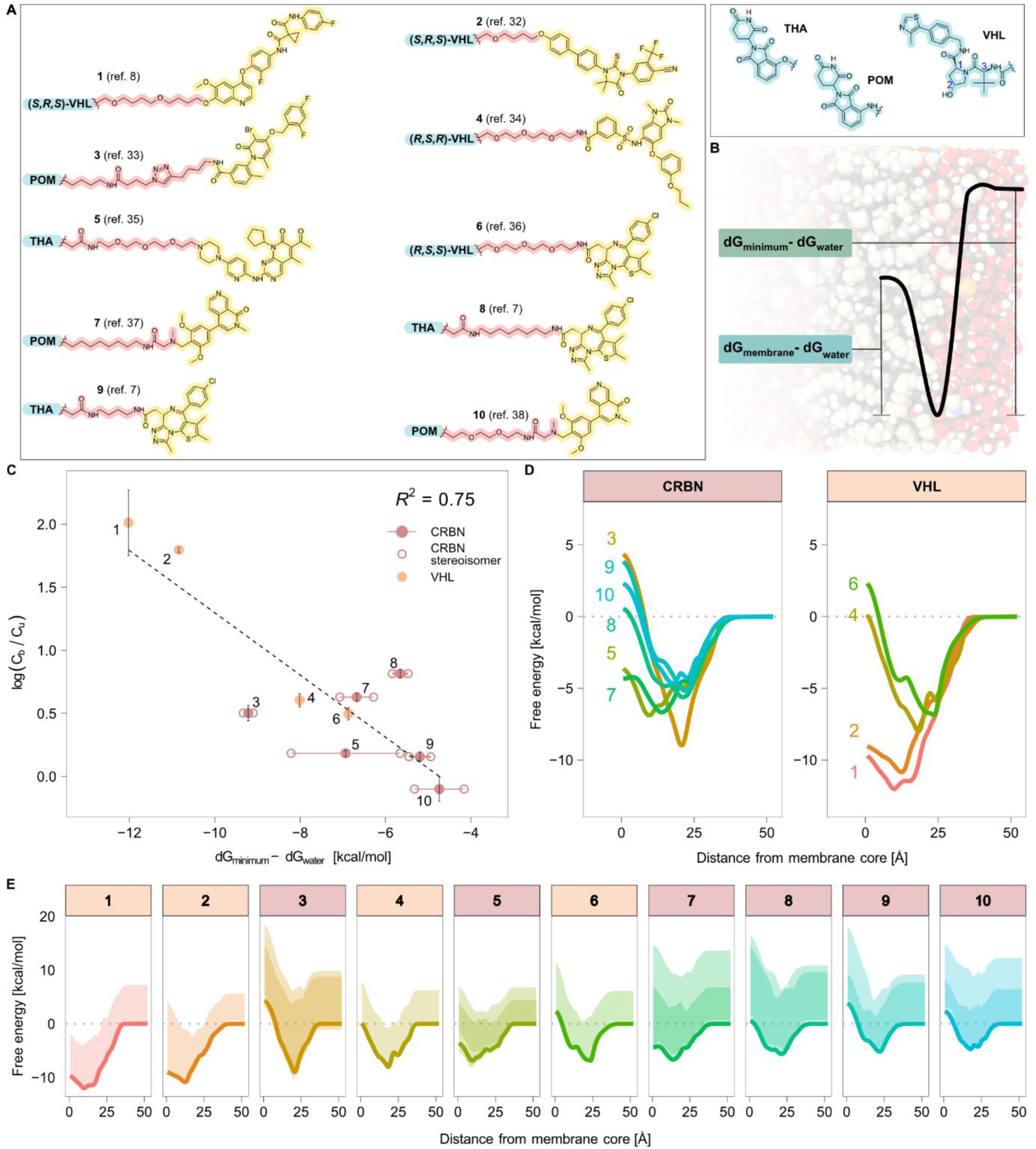
Interactions of PROTACs with *in vitro* and simulated cell membranes. A) PROTAC molecular structures^7,8,32–38^ B) Schematic illustration of a typical free energy profile in a simulated DMPC membrane with calculated energy barriers. C) The depth of energy minima correlates well with experimental binding to HEK293 cell homogenate, expressed as the logarithm of bound-to-unbound concentration ratios, log (C_b_/C_u_) (mean ± SD, n = 3). Full circles show the average and open circles show the individual stereoisomers. D) Average free energy profiles for CRBN-binding PROTACs (left panel) and VHL-binding PROTACs (right panel). E) Free energy profiles of individual PROTACs with conformational variability. Solid lines indicate the Boltzmann-weighted energy profiles, averaged across molecule stereoisomers. Shaded areas show the min-max range of free energies across all conformations for each individual stereoisomer. Darker shades indicate energy regions covered by both stereoisomers.

Longer alkane linkers resulted in higher degree of binding (4.6-fold higher bound-to-unbound concentration ratios for compound **8** than **9**), and a PEG based linker resulted in lower membrane binding than an alkane based one of comparable length (5.3-fold lower for compound **10** than for **7**). Thus, it is clear that the choice of linker can substantially influence interactions with cellular membranes.

The test compounds were further analyzed *in silico* with the aim to quantitate the energetics of membrane interactions and identify the main barriers and preferential orientations and conformations adopted in the membrane binding. We used the continuum solvation model (COSMO) in which density functional theory (DFT) based calculations are applied to derive surface polarization charge densities for each type of molecule in a multi-component mixture, and their miscibility subsequently assessed through statistical thermodynamics. To assess conformation-dependent effects, charge densities were calculated for representative low-energy conformations of each analyzed PROTAC obtained from molecular mechanics calculations, and a Boltzmann weighting approach was used to derive free energy profiles for the interaction of each compound’s conformational ensemble with a DMPC (1,2-Dimyristoyl-sn-glycero-3-phosphocholine) phospholipid membrane (**Fig. 3D-E**). We found that deeper energy minima strongly correlated with increased binding to cell homogenate (**Fig. 3C**), in line with similar calculations for small organic solutes where COSMO-derived energy minima have been shown to reflect affinity to model phospholipid membranes. ^30^ VHL-binding PROTACs were, on average, more bound to cell homogenate and had correspondingly more pronounced energy minima than CRBN-binding ones.

Encouraged by the correlation between experimentally measured affinities for isolated cell membranes and COSMO-derived free energy profiles, we extended our analysis to a more comprehensive set of 157 representative molecules to probe in detail what structural and chemical features govern the respective profile shapes and energy barriers.

Increased molecule lipophilicity (assessed with the two-dimensional cLogP estimate), on average, led to deeper energy minima at the water–lipid interface and lower barriers for traversing the membrane core (**Fig. 4A**). Weaker, positive, correlations were seen between energy barrier height and the polar surface area exposed in the most shielded conformation, indicating that the ability to form such low-polarity conformations are beneficial for interactions in the membrane core (**Fig. 4B**). No apparent correlations were seen for the other fundamental molecular properties assessed (MW, HBD, HBA, TPSA and nRotB).

**Figure 4.**
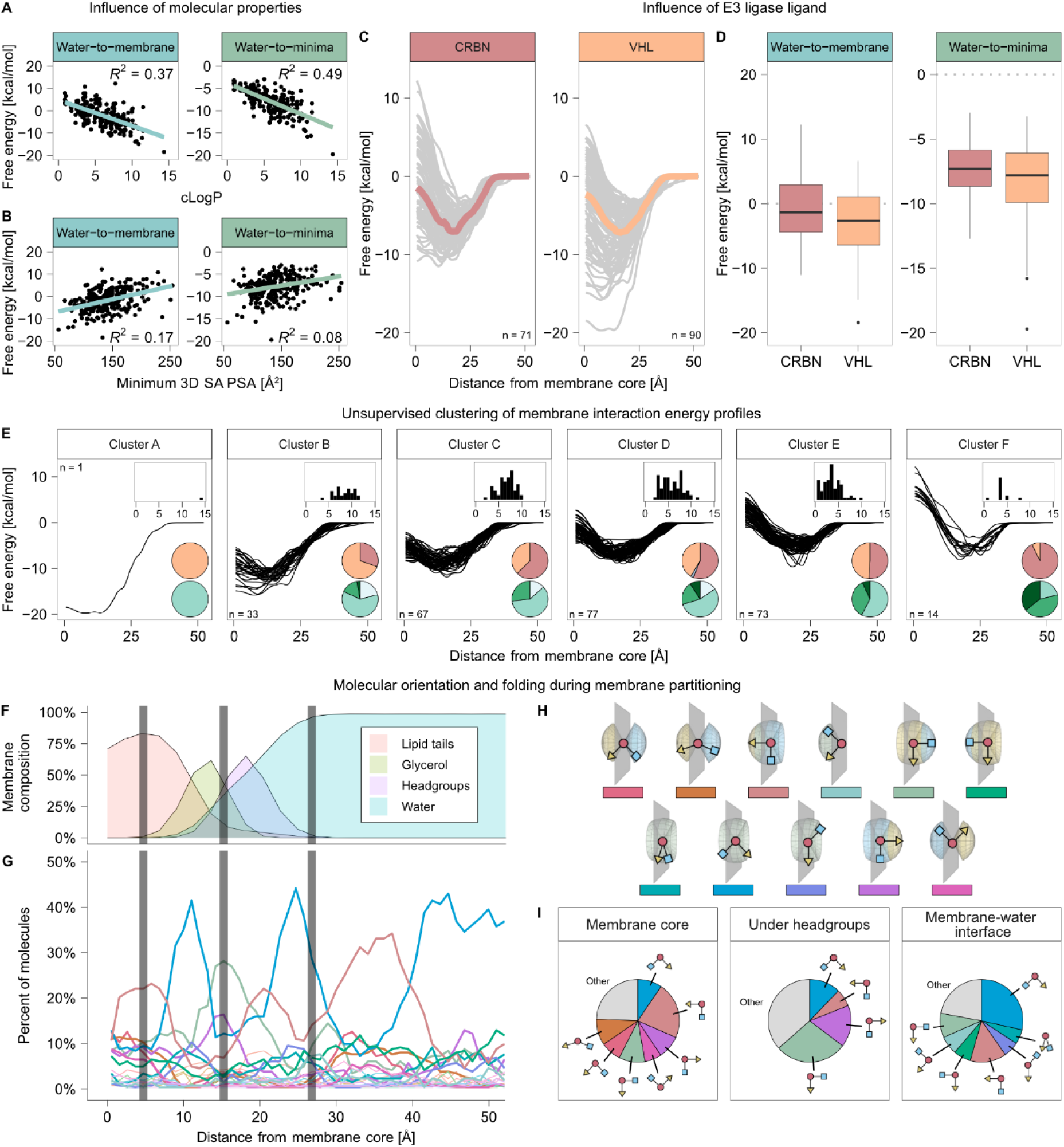
Interactions of representative PROTACs (n=157) with simulated phospholipid membranes. A) High cLogP correlates with lower energy minima and barrier in the membrane core. B) Lower minimum solvent accessible polar surface area correlates moderately with lower energy in the membrane core. C) Free energy profiles of CRBN- and VHL-binding PROTACs. Median profile in red/orange, generated from the median energy at each depth in the membrane. Per-compound profiles in grey. D) Free energy barriers of CRBN- and VHL-binding PROTACs at membrane core and minima. Black solid line indicates the median and the bottom and upper box limits are the 25^th^ and 75^th^ percentile. Whiskers extend to 1.5 multiples of the interquartile length (IQR) and dots represent observations that extend outside of 1.5 IQR. Energy barrier explanation in figure legend 4B. E) Unsupervised clustering of free energy profiles, cLogP distributions are indicated in histogram inserts. Top pie charts show proportion of VHL-binding (orange), CRBN-binding (red) and IAP-binding (grey) PROTACs. Bottom pie charts show minimum 3D SA PSA; < 100 Å^2^ (white), 100-150 Å^2^ (light green), 150-200 Å^2^ (green) or > 200 Å^2^ (dark green). F) Membrane composition. G) Orientations of PROTACs at different depths of the membrane. Colors represent different shapes and orientations of PROTACs relative to the membrane, as indicated in H. H) Schematic PROTAC orientations. Grey area indicates membrane slice. I) Molecule orientation at membrane core, under the headgroups and at the membrane-water interface.

Consistent with the experimental dataset (**1**–**10**), energy minima at the interface were generally lower for VHL-than for CRBN-directed PROTACs, and the energy barrier at the membrane core were also lower for VHL-directed PROTACs (**Fig. 4C-D**). In contrast, molecular properties of the POI ligand moieties did not noticeably correlate with the sizes of energy minima or barriers, except for a weak correlation between the former and the cLogP of the POI ligand (**Supplementary Fig. 3**).

Using unsupervised clustering to identify groups of PROTACs that share similar energy profiles further confirmed the impact of molecule lipophilicity (**Fig. 4E**). Furthermore, the proportion of CRBN-binding PROTACs was greater in clusters with more pronounced membrane penetration barriers, and PROTACs with lower solvent-accessible polar surface area were more common in clusters with small penetration barriers. Linker length and chemistry had less clear-cut effects on PROTAC–membrane interaction energies. Longer PEG-based linkers were more common in clusters with greater penetration barriers (clusters D-F), whereas rigid and lipophilic linkers (ring- and alkyl-based linkers) were more common in the low-barrier clusters A-B. (**Supplementary Fig. 4**). Longer alkyl-based linkers yielded, on average, lower energy minima than other linker classes (**Supplementary Fig. 5**).

The observed environment-dependent exposure of polar atoms (**Fig. 2A and 2D**) indicated functionally selective conformational sampling as the PROTACs partition into the hydrophobic environment of the membrane core, but the implicit-solvent calculations used are not sufficient to reveal how the permeating molecules orient themselves as they enter the membrane. We therefore extended our analysis, examining the preferential orientation of PROTACs as a function of penetration depth in the heterogeneous QM-derived membrane model (**Fig. 4F-I**).

At the water–membrane interface the PROTACs tended to orient themselves flat against the membrane surface, while when positioned immediately beneath the headgroups the E3 ligand more commonly pointed out towards the water–in particular for CRBN-directed PROTACs. Within the lipid tail region of the membrane, the dominating orientation was with the POI ligand pointing towards the core. This is reasonable, given that the POI ligand is the most lipophilic part of the PROTAC and thus expected to interact the most with the phospholipid tails. In the clusters of similar free energy profiles (**Fig. 4E**), compounds with low penetration barriers at the membrane core (clusters A+B) more often exhibited stretched conformations, with the POI ligand pointing towards the membrane center and the E3 ligand pointing towards the water, whereas in clusters with higher membrane barriers (clusters D-F) the E3 ligand more commonly pointed inwards to the membrane center (**Supplementary Fig. 6**). Interestingly, when the PROTAC centers-of-mass were positioned in bulk water at a distance of ∼35 Å from the membrane core, the dominating orientation was with the POI ligand towards the membrane core, suggesting a preference for initiating interactions with the phospholipid headgroups through their POI ligand domain.

Assessing exposed surface polarity for the conformational ensembles at different depths in the model lipid membrane revealed a more diverse picture. While the majority (54%) of the studied molecules predominantly sampled lower-polarity conformations in the membrane core than in water (with up to 102 Å^2^ reduction in exposed PSA; **Supplementary Table 3**), a substantial proportion of the compounds showed the opposite behavior, with 11% of all compounds exposing 10 Å^2^ or more excess PSA in the hydrophobic membrane environment than in water. Notably however, almost all (89%) of the molecules exposing more polar surface in the membrane core also had a correspondingly increased *total* surface area. This indicates that such molecules adopt more stretched conformations in the membrane core, exposing more polarity, but at the same time also exposing more non-polar atoms, compensating for the desolvation penalty.

### Membrane affinities in defined structure series

Our global analyses thus indicated that PROTAC–membrane interactions are influenced by both E3 ligand and linker properties, suggesting opportunities to optimize PROTACs for a given protein of interest by modifying these functional units. To confirm these findings and assess membrane affinity-driving features more systematically, we designed and synthesized a series of new PROTAC molecules, incorporating identical POI ligands but with variations in E3 ligand moiety and linker length and chemistry (**Fig. 5A**) and measured their membrane affinities experimentally.

**Figure 5.**
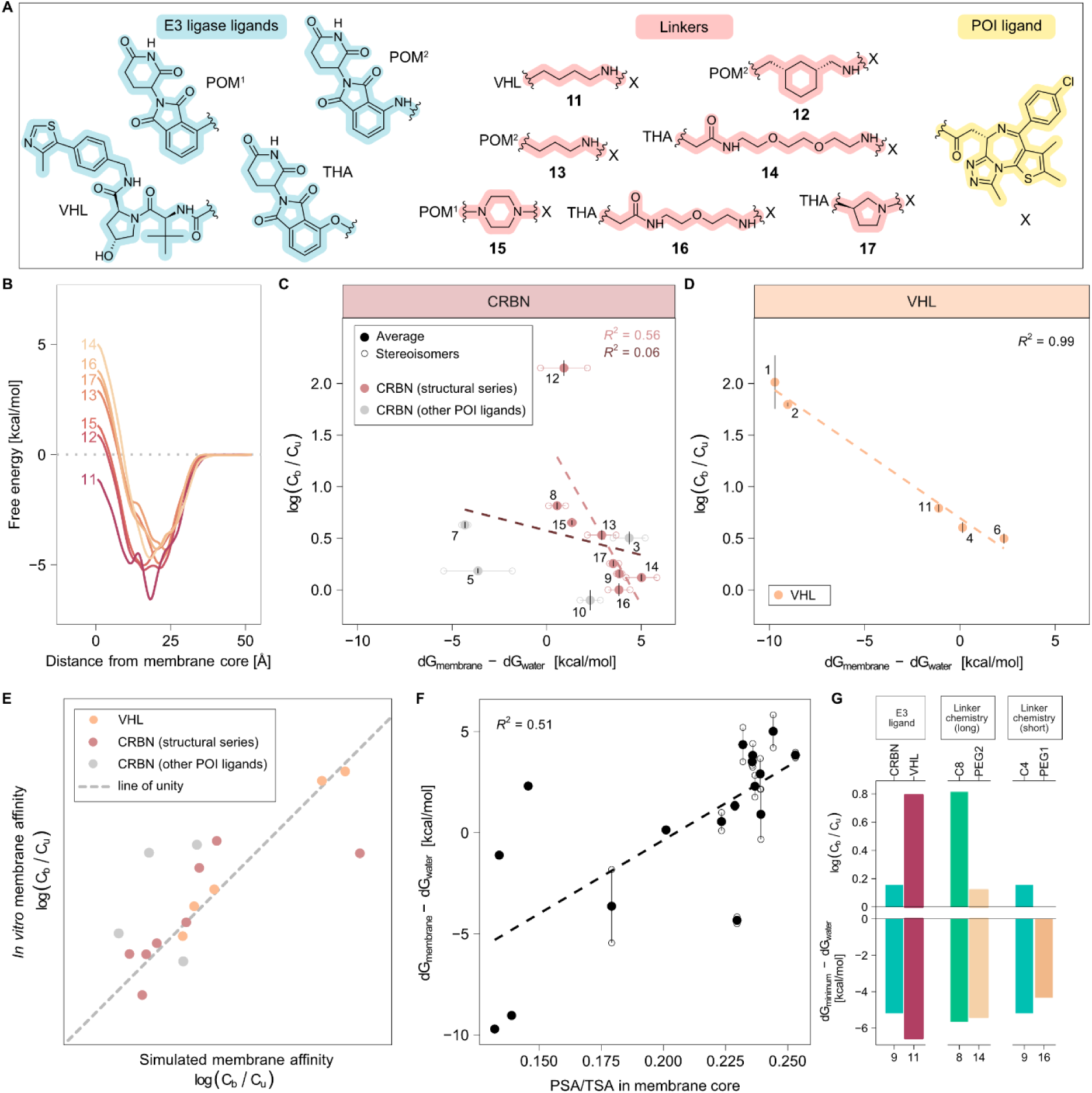
Cell membrane affinities of a new series of structurally related PROTACs. A) Molecular structures of PROTACs **11**-**17**. B) Average free energy profiles for compounds **11-17**. C-D) Bound-to-unbound concentration ratios (mean ± SD, n = 3) correlate strongly to the water-to-membrane core energy barrier for VHL-binding PROTACs (panel D, orange full circles), while a more moderate correlation is seen for CRBN-binding PROTACs (panel C) that share the same E3 ligand and POI ligand. Full circles show the average of a 1:1 racemic mixture and open circles show values for the individual stereoisomers. PROTACs highlighted in red belong to a structural series with identical E3 ligand and POI ligand while grey circles show additional CRBN-binding PROTACs that are more structurally diverse. E) Correspondence between simulated and *in vitro* measured membrane affinities. F) Energy barrier water-to-membrane core correlates moderately to fraction solvent exposed polar surface area of total solvent exposed surface area. Full circles show the average of a 1:1 racemic mixture and open circles show values for the individual stereoisomers. G) Comparison of energy barriers and experimental membrane binding. Top: Bound-to-unbound concentration ratios (mean). Bottom: Depth of energy minima. Left: The VHL-binding **11** has a lower energy minimum and higher cell homogenate binding than the CRBN-binding **9**. Middle: **8** with a long alkane linker has a lower energy minimum and higher experimental binding than **14** with an equally long PEG-linker. Right: **9**, with a shorter alkane linker has a lower energy minimum and higher cell homogenate binding than **16** with a PEG-based linker.

This expanded set confirmed the initial trend with observations of increasing membrane partitioning for PROTACs with lower energy minimum at the water–membrane interface and greater molecular lipophilicity (**Supplementary Fig. 7**). Exchanging the E3 ligand resulted in a 3-fold higher bound-to-unbound ratio of the VHL-directed **11** than the linker- and POI-matched CRBN-directed **9**, and a corresponding deeper energy minimum for **11**. Thus, the substitution of E3 ligase ligand strongly affects membrane affinity, most likely due to the difference in lipophilicity that follows. We also confirmed our previous findings that linker length and chemistry affect membrane partitioning and depth of the energy minima, with PROTACs containing alkane linkers exhibiting higher binding and deeper energy minima than structurally matched compounds containing equally long PEG-based linkers. (**Fig. 5G**).

Despite relatively small differences in the overall molecular structure, the new series of analogs varied widely in the height of the energy barrier at the membrane core—similar to observations for compounds **1**–**10** (**Fig. 3D and 5B**). This energy barrier is associated with both membrane affinity and the rate of solute permeation (flip-flop) across the lipid bilayer. ^31^

Notably, there was a clear correlation between barrier height and *in vitro* membrane affinity for the VHL-directed PROTACs in our set. A similar trend was seen for the subset of CRBN-directed PROTACs that share the same POI ligand, whereas global trends in the full set of CRBN-directed PROTACs were obscured by the wide variation in POI ligand properties (**Fig. 5C-E**). Interestingly, throughout the VHL series the ligase ligand remained identical and all compounds but one (compound **11**) contained similar PEG-based linkers, whereas in the CRBN analogs structure variations arise solely from the linker domain and the nitrogen versus oxygen connecting atom in pomalidomide/thalidomide-based E3 ligands. Thus, PROTAC– membrane affinities can be modulated by isolated variations in either the POI ligand or the linker regions, indicating that rational optimization of the linker region can be used to balance properties in the functionally active domains.

Analyzing the preferential orientation of the experimentally tested PROTACs, there was a clear tendency to orient the POI ligand towards the membrane center and the ligase domain either towards the water phase or along the membrane surface. This is likely explained by a greater relative lipophilicity of the POI ligand. The trend is more pronounced in the CRBN PROTACs, consistent with the greater difference in lipophilicity between the ligase and POI binding domains in these compounds.

Finally, compounds with proportionally greater exposed polar surface in general exhibited higher barriers at the membrane core (**Fig. 5F**). The trend was more evident in CRBN-binding than VHL-binding PROTACs, indicating that the ability to shield polarity is more important for the former and consistent with the less lipophilic nature of the CRBN ligands (**Table 1**). The E3 ligand of CRBN-binding PROTACs also more often pointed out towards the water, indicating a preference for interactions with the lipid headgroups. Flip-flop across the membrane can thus be assumed to be more difficult for this class of PROTACs, both as a result of an increased desolvation cost at the interface, and a higher barrier across the membrane core.

## Discussion

Through systematic analyses we provide insights into how the E3 ligand, the linker chemistry, and general molecular properties affect the affinity of PROTACs for cell membranes and ultimately their cell permeability and intracellular exposure.

Previous analyses of conformational dynamics of linear and macrocyclic beyond-Rule-of-5 drugs have indicated that polarity shielding is a major factor in determining cell permeability. ^18–22^ We see similar global trends in PROTACs, with lower exposed polar surface in non-polar than in polar environments, based on conformations sampled using molecular mechanics.

Further, using quantum mechanics-based calculations to delineate PROTAC interactions with simulated lipid membranes, we see that a majority of all compounds analyzed sample conformations with lower exposed polarity in the membrane core than in the water phase. At the same time many PROTACs, when located in the core of the lipid membrane, preferentially orient in extended conformations pointing along the membrane normal, which in several cases led to *greater* average exposed polarity in the membrane core than in water. That said, compounds with such behavior typically also exposed a higher *total* surface area, and thus that the energy expense of exposing polar atoms in a non-polar environment is compensated by an increased surface hydrophobicity.

An increase in the conformation-independent property cLogP, as well as experimental values of cell homogenate binding was associated with deeper energy minima at the membrane– water interface, while lower solvent exposed polar surface area correlated with a decreased energy barrier in the transition from the energy minima to the membrane core, albeit more weakly. These results suggest that the interactions of PROTACs with the lipid membrane are driven largely by the overall molecular lipophilicity, and is compensated by the ability to shield polar functionality. Such environment-dependent polarity shielding appears more pronounced for CRBN-binding than for VHL-binding PROTACs, and may partly compensate for the more polar nature of the CRBN ligands.

Notably, conformation-dependent shielding of polar surface area correlated much more strongly to the overall folding of the molecule, rather than the folding of the linker region alone. Thus, conformational rearrangement within the POI ligand and/or E3 ligand domains contribute to the observed polarity shielding. We note that there are clear differences between different linker types in the propensity of the PROTACs to form folded structures, which suggests that the choice of linker plays an important role in the overall molecule folding.

Long PEG-based linkers–with or without ring systems in the linker–resulted in the most folded molecules and are thus assumed to reduce exposed polar surface area the most. The caveat to this assumption is the greater polarity of the PEG linkers themselves, which tend to remain solvent-exposed also in folded structures.

Finally, different membrane interactions were observed for VHL- and CRBN-binding PROTACs, where the former in general have smaller energy barriers in the membrane core, which is typically associated with a more favorable membrane permeability. ^31^ The greater lipophilicity of VHL-binding PROTACs may, however, lead to a more problematic solubility than CRBN-binding PROTACs.

In summary, an intricate picture emerges in which conformational flexibility is beneficial for PROTAC–cell membrane interactions and cell permeability by enabling environment-dependent shielding of polarity, but is contrasted with the preference for extended conformations within the membrane. It thus appears that shorter-range intramolecular interactions leading to polarity shielding selectively within the POI ligand and/or E3 ligand domains may be preferred over more drastic molecular folding, in order to design PROTACs with improved interactions with cell membranes.

## Methods

### Calculation of 2D descriptors

SMILES codes from the original data sources or manually sketched molecules were converted to canonical format using OpenBabelGUI (v 3.1.1). ^39^ E3 ligands were identified through visual inspection of the molecules, as were the POI ligands. All molecules were then decomposed into their constituent E3 ligand, linker and POI ligand domains using a custom RDKit (v 2021.09.2) script implemented in Python (v 3.9.7). ^40,41^ 2D descriptors, including molecular weight (MW), calculated LogP (cLogP), hydrogen bond acceptor and donor counts (HBA / HBD), number of freely rotatable bonds (nRotB), and topological polar surface area (TPSA) were calculated using RDKit for the intact molecule as well as for the different structural domains.

The most common linker patterns were identified and matched to all molecules in the database using RDKit. The classification was visually verified and linker fragments that had been incorrectly mapped (e.g., falsely assigned to the butane group when it contained a longer alkyl segment) were manually reassigned. Further, the identified linkers were grouped together with chemically similar linkers, to allow analysis of how linker chemistry affects membrane interactions. First, all linkers were divided into two groups; those containing rings and those with only linear chains. Then, depending on length of linear chains (with or without ring systems attached), linkers were divided in Short (two to eight atoms) or Long (nine atoms and longer). Non-ring linkers were separated based on linker chemistry into subgroups (Short/Long Alkane or Short/Long PEG). Ring-containing linkers were separated into subgroups of Only Ring (no linear chains attached), Short Ring (two to eight atoms linear chain of either alkane or PEG chemistry) and Long Ring (9 or more atoms of either alkane or PEG chemistry). Linkers which did not fit into any, or in more than one of the groups, were placed in a separate group (Others).

### Conformational sampling

3D starting geometries were prepared from SMILES strings in LigPrep (Schrödinger Release 2021-3: LigPrep, Schrödinger, LLC, New York, NY, 2021) using the OPLS4 forcefield. ^42^ Molecules were desalted, tautomers were generated and previously specified chiralities were retained. Stereoisomery of non-defined stereocentra were varied for all molecules and ionization states relevant at a pH of 7.0 +-0.2 were generated for 400 of the 891 molecules. An unionized state was always generated.

Conformational sampling of the generated variants was performed in water and chloroform, using MacroModel (Schrödinger Release 2021-3: MacroModel, Schrödinger, LLC, New York, NY, 2021) and the OPLS4 forcefield. Default settings were applied if not otherwise stated. Minimization was performed using the Polak-Ribier Conjugate Gradient (PRCG) method using a maximum of 5000 iterations per molecule. Minimizations were set to converge on gradient with a threshold of 0.05. The conformational sampling used the Mixed torsional/Low-mode sampling method with automatic setup. An intermediate torsion sampling option was used. Mirror-image conformations were retained and the maximum number of steps was set to 5000, with 100 steps per bond. The energy window for saving structures was set to 21.0 kJ/mol and redundant conformers were eliminated using a maximum atom deviation of 0.5 Å. Molecule variants containing negatively charged sulfonamides were not possible to process in the chloroform environment, and were thus excluded from further analysis.

### Calculation of 3D descriptors

A custom PyMol (v 2.1) script^20^ was used to calculate van der Waals and solvent-accessible (rolling water probe) polar and total surface areas (PSA/TSA) of each of the conformations generated in the conformational sampling.

Conformational flexibility in the linker region and in the full molecule were calculated by identifying the centers of mass of the E3 ligand and POI ligand domains, as well as the anchor points for the linker, in both 2D and 3D structures. The 2D structures used were generated in RDKit from SMILES strings and the 3D structures were the conformers generated in the conformational sampling.

The maximum distance between E3 ligand and POI ligand center of masses (representing the distance in a fully stretched molecule) was calculated from the 2D structures, by taking the sum of the distances between E3 ligand center of mass and E3 ligand-linker anchor point (d_E3 ligand, 2D_), E3 ligand linker anchor point and POI ligand linker anchor point (d_linker, 2D_), and POI ligand linker anchor point and POI ligand center of mass (d_POI ligand, 2D_) (Eq. 1).

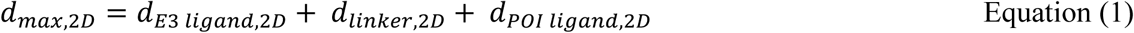

The distance between E3 ligand center of mass and POI ligand center of mass (d_E3 ligand to POI ligand, 3D_) was calculated for each 3D conformer and compared to the maximum distance, giving the measure *‘molecule contraction’* (Eq. 2).

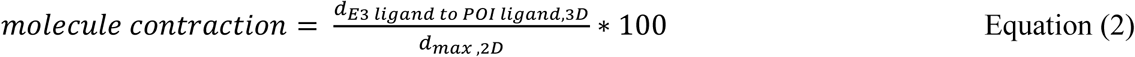

Analogously, the degree of linker stretch was calculated for each 3D conformer and compared to the maximum linker length (‘*linker contraction’*) (Eq. 3).

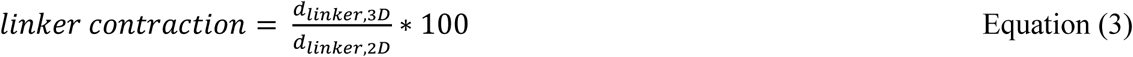

### Molecular dynamic simulation of DMPC membrane model

A membrane model was simulated, consisting of DMPC lipids. The system was simulated at 303 K, using 70 water molecules per lipid. The atomistic molecular dynamics (MD) simulations were performed in the GROMACS software package (version 2021.3). ^43–49^

A pre-equilibrated DMPC bilayer was obtained, ^50,51^ containing 128 DMPC lipid molecules as well as water. The TIP3p water model was used according to literature recommendations. ^52^ To ensure enough room for the PROTACs in the solvent phase, the box size was adjusted in the z dimension from 62 Å to 108 Å and additional solvent was added with a concentration of 0.15M NaCl using GROMACS and a perl script (http://www.mdtutorials.com/gmx/membrane_protein/Files/water_deletor.pl).

Energy minimization of the lipid membrane was performed using a steepest descent algorithm with 50’000 steps. Long-range electrostatic interactions were calculated using the particle-mesh Ewald method^53^ with a cut-off of 1.4 nm. Van der Waals interactions were treated with a plain cut-off at 1.4 nm. ^54^ Long-range dispersion corrections for energy and pressure were added. ^55^ Periodic boundary conditions were imposed in every dimension.

The simulation was run in semi-isotropic conditions where pressure in the xy plane was coupled separately from the z direction. Pressure was kept constant at 1 bar using the Parrinello-Rahman barostat, ^56^ with a coupling constant of 10.0 ps and a compressibility of 4.5 x 10^-5^ bar^-1^. The temperature was kept constant at 303 K using the Nosé-Hoover thermostat, ^57,58^ coupling the membrane and solvent separately, with a coupling constant of 0.5 ps. The P-LINCS algorithm was used to constrain all covalent bonds in the lipids^59^ and for water, the SETTLE method was applied, ^60,61^ allowing for a timestep of 2 fs, enforced using the leapfrog integrator. ^62^

The membrane was equilibrated in two steps, first with an NVT (Number of particles, Volume and Temperature) ensemble for 10ns with a 2 fs timestep, followed by an NPT (Number of particles, Pressure and Temperature) ensemble for 40 ns. Pressure, temperature, density and box vectors were verified to have stabilized around desired values, with no trends. Production simulations were run for 100 ns with a timestep of 2 fs. Coordinates were saved every 1 ps and the neighbor list was updated every 10th step.

### COSMO calculations

In COSMOtherm simulations, statistical thermodynamics are applied to surface polarization charge densities obtained from density functional theory (DFT) optimizations^63^ separately performed for each component of the simulation, i.e. water, sodium cation, chloride anion, a representative DMPC lipid and a PROTAC molecule in its unionized state. A representative lipid conformation was selected in a multi-step process. First, the lipid membrane MD frame with the area per lipid most closely representing the average was selected. All 128 lipids in this frame were subjected to DFT/COSMO calculations in COSMOconf. From these, the RMSD compared to the conformers observed in the MD simulation was calculated using GROMACs tool *gmx rms*, using the trajectory from the MD simulations as reference. The most representative (lowest RMSD) low-energy conformation was selected for further calculations in COSMOtherm. Notably, COSMOtherm is robust to the choice of lipid conformer, as long as it is fairly representative. ^64^ The BP-TZVP basis set was used and DFT optimizations were performed in COSMOconf (Biovia COSMOlogic v 21.0) to obtain sigma profiles.

Six frames from the equilibrated DMPC membrane were separately imported into COSMOtherm (Biovia COSMOlogic v 21.0) as a planar membrane, and divided into 50 slices of approximately 0.1 nm thickness, as recommended in literature. ^64^ Each slice was treated as a homogenous phase with its own sigma profile. Input files for water, Na cation, Cl anion were used as supplied with the COSMO software suite, and were added to the membrane file together with the selected DMPC lipid conformer. Unionized PROTAC molecules were then added to the system and rotated in 162 different orientations in each layer using the COSMOtherm plugin COSMOmic to generate layer-dependent solvation energies with the BP_TZVP_21 parametrization. Due to the size of PROTAC molecules they are likely to span over several layers. Parts that extend into adjacent layers were fitted to those respective layers. Local sigma profiles were calculated for each layer and an integral sigma profile was generated. From this, the solvation energy for each molecule in each layer was calculated and a free energy profile was generated.

### Hierarchical clustering

To identify molecules with similar membrane interaction behavior, free energy profiles generated from the combined conformer sets were hierarchically clustered using the hclust function with the complete linkage method in R Stats Package in R (v.4.2.2). ^65^

### Calculation of conformational ensemble

To elucidate the conformation-dependent free energy of solvation of the studied PROTACs at different positions in a phospholipid membrane, free energy profiles from combined conformer sets for each molecule, as well as for their individual conformers, were calculated at 310 K in DMPC membranes using COSMOtherm. ^63^ Free energy calculations were repeated using each of six snapshots captured over the MD simulation timeframe. To allow averaging the free energy profiles at identical positions along the membrane normal, the profiles were linearly interpolated using R, and then average free energy profiles were calculated for each conformer set or conformer. The profiles for the separated conformers were Boltzmann weighted using a custom R script to derive the highest-probability conformational ensemble in each section of the membrane.

### Calculation of molecule orientation

To assess the preferential orientation and bending of PROTAC molecules at various positions in the DMPC/water system, the molecules were separated in their E3 ligand, linker and POI ligand domains and the relative positions of the respective weighted centers of mass (hydrogens excluded) were determined. The molecule was kept at a fixed orientation in COSMOtherm and the membrane was rotated to find the lowest energy orientation. To allow comparison among conformations, the coordinate systems were first realigned. The most probable orientation of each molecule was then identified at different distances from the membrane core. First, the linker center of mass was aligned to origo. Then, three separate angles were calculated; 1) between the membrane normal and linker to E3 ligand vector, 2) between the membrane normal and the linker to POI ligand vector and 3) between the linker to E3 ligand vector and linker to POI ligand vector. Each type of angle could take a value of 0° to 180° and was split in three 60° sections. This generated 27 different groups to which each molecule was assigned at every section of the membrane.

### Determination of fraction of unbound drug in cell homogenate

Reagents and cell culture media were obtained from Sigma-Aldrich (St Louis, MO) or Invitrogen (Carlsbad, CA). Commercially available PROTACs were obtained from Tocris (Bristol, UK), new PROTACs were synthesized by SciLifeLab (Solna, Sweden) and small molecule drugs were obtained from Sigma-Aldrich (St Louis, MO) and were of ≥ 95% purity. Stock solutions of compounds (10 mM) were prepared in dimethyl sulfoxide (DMSO) and stored aliquoted at -80 °C. Reagent tubes and plates were Eppendorf Protein LoBind (Hamburg, Germany).

The unbound fraction of a selected series of PROTACs was determined through equilibrium dialysis against cell homogenate. HEK293 cells (ATCC, Manassas, VA) were grown in Dulbecco’s modified Eagle’s medium (DMEM) supplemented with 10% fetal bovine serum (FBS) at 37 °C, 95% relative humidity and 5% CO_2_. Cell pellets were harvested using 0.05% trypsin and centrifuged at 260 xg for 5 min. The cells were resuspended in Dulbecco’s phosphate buffer (DPBS), counted using a Luna II cell counter (Logos Biosystems, Anyang, South Korea) and centrifuged at 260 xg. The supernatant was discarded, and cell pellets stored at -80 °C prior to cell homogenate preparation. On the day of the experiment, cell pellets were thawed, resuspended in Hank’s balanced salt solution (HBSS) buffered with 18.4 µM HEPES to a concentration of 10×10^6^ cells/ml. The cell suspension was homogenized using ultrasonication for 10s at 20% intensity while kept on ice.

Binding to cell homogenate was determined at 0.1 µM for all compounds. A Rapid Equilibrium Dialysis device (Thermo Fisher, Waltham, Massachusetts) was used to dialyze the samples by adding 200 µl compound spiked cell homogenate to one chamber and 350 µl HBSS/HEPES to the other chamber. The cassette was incubated on an Eppendorf ThermoMixer C (Hamburg, Germany) at 900 rpm for 4 hours at 37 °C.

Compound stability was assessed by adding compound-spiked cell homogenate to an empty well in the dialysis cassette. At the end of the experiment these concentrations were compared to identical control samples stored at 4 °C. Mass balance was calculated at the end of each experiment. Further, in all wells two control compounds (atorvastatin and lopinavir) were included to ensure that the assay plate was functional. All compounds were tested in triplicate in one experiment. Compound **2** was only detected on the cell homogenate side, and thus the concentration on the buffer side was set to half of the lower limit of quantification to allow calculation of an approximate cell homogenate binding.

Final samples were matrix matched by taking 1 part sample and 1 part blank cell homogenate or buffer. Proteins were precipitated using 1 part matrix matched sample and 3 parts acetonitrile supplemented with 0.3% formic acid (FA) and 5 nM verapamil (internal standard), and this was diluted 1:1 with MilliQ prior to centrifugation at 2500 xg for 20 min. Compound concentrations in the supernatant were quantified using a reversed phase system with a Waters Acquity UPLC HSS T3 column (50 mm×2.1 mm, 1.8 μm) in an Aquity UPLC connected to Xevo TQ-XS mass spectrometer (Waters Corp., Milford, MA). The chromatographic run was done using solvent A (0.1% FA in MilliQ) and solvent B (0.1% FA in acetonitrile Optima LC/MS grade (Fisher Scientific) with isocratic conditions between 0-0.3 min (99.8% solvent A and 0.2% solvent B), a linear gradient between 0.3-1.3 min (from 0.2% to 95% solvent B), isocratic conditions between 1.3-1.8 min (95% solvent B) and equilibration back to start conditions between 1.8-2.3 min. Mass transitions, cone voltages and collision energies are found in **Supplementary Table 4**.

### General chemistry methods

All solvents and reagents were used as received from commercial suppliers. Column chromatography was employed on normal-phase silica gel (40-60 µm, 60 Å). ^1^H-nuclear magnetic resonance (NMR) and ^13^C-NMR spectra were recorded on a 400 MHz spectrometer at 298 K and calibrated using the residual peak of the solvent as an internal standard: CDCl_3_ (δ_H_ 7.26 ppm, δ_C_ 77.16 ppm), CD_3_OD-*d4* (δ_H_ 3.31 ppm, δ_C_ 49.00 ppm), DMSO-*d6* (δ_H_ 2.50 ppm, δ_C_ 39.52 ppm). NMR spectra are reported as follows: chemical shift, multiplicity (s = singlet, d = doublet, t = triplet, q = quartet, m = multiplet, dd = doublet of doublets), coupling constant (*J*) in Hertz (Hz) (if applicable) and integration (proton spectra only). NMR spectra are found in Supplementary Information. Analytical liquid chromatography coupled with mass spectrometry (LCMS) was performed using a column ACE 3 C8 (50 × 3.0 mm); water (0.1% TFA) and acetonitrile were used as mobile phases at a flow rate of 1 mL/min, with a gradient time of 3.0 min. High-resolution mass spectra were recorded on a QTOF mass spectrometer with electrospray ionization (VIP-HESI). Ions were detected in positive mode. ESI-L Low Concentration Tuning Mix was used for instrument calibration prior to measurements. Samples were prepared as 1 µM solutions in ACN/water 1:1 v/v and directly infused into the mass spectrometer at a constant rate of 10 µL/min.

Synthesis of (2S,4R)-1-((2S)-2-(5-(2-((6S)-4-(4-chlorophenyl)-2,3,9-trimethyl-6H-thieno[3,2-f][1,2,4]triazolo[4,3-a][1,4]diazepin-6-yl)acetamido)pentanamido)-3,3-dimethylbutanoyl)-4-hydroxy-N-(4-(4-methylthiazol-5-yl)benzyl)pyrrolidine-2-carboxamide (**11**).

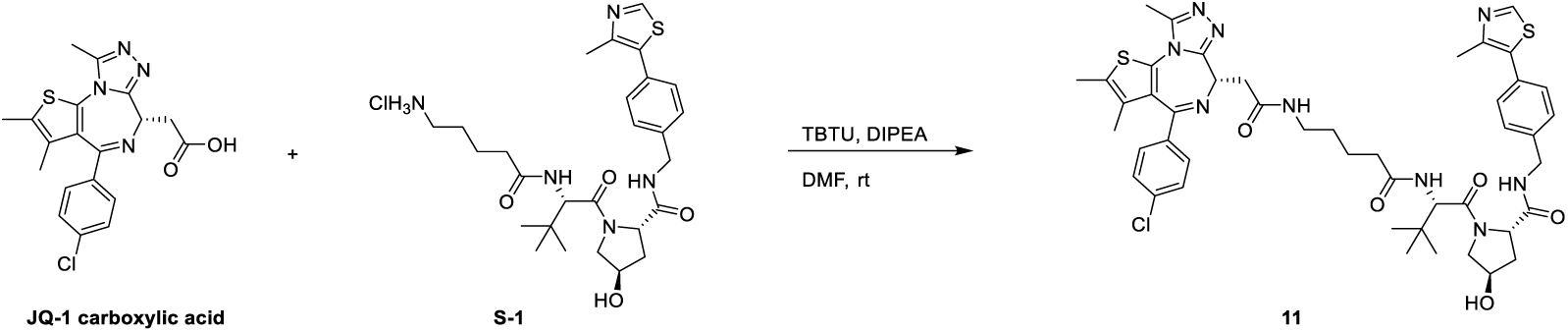

DIPEA (17 µL, 0.1 mmol) was added to a stirred solution of JQ-1 carboxylic acid (10 mg, 0.025 mmol) and TBTU (8.4 mg, 0.026 mmol) in DMF (0.5 mL). After 30 minutes, **S-1** (13 mg, 0.025 mmol) was added and the reaction was allowed to perform for 2h. The mixture was diluted with EtOAc, washed with NaHCO_3_ sat. aq., dried over sodium sulfate, filtered and concentrated to dryness. The resulting material was purified using automated flash chromatography (12 g silica cartridge, DCM/MeOH: 0-10 % over 7 column volumes) to provide compound **11** (16 mg, 70 % yield). ^1^H NMR (400 MHz, DMSO-*d6*) δ 8.98 (s, 1H), 8.56 (t, *J* = 6.1 Hz, 1H), 8.18 (t, *J* = 5.5 Hz, 1H), 7.88 (d, *J* = 9.4 Hz, 1H), 7.57 – 7.34 (m, 8H), 5.13 (d, *J* = 3.6 Hz, 1H), 4.55 (d, *J* = 9.5 Hz, 1H), 4.50 (dd, *J* = 8.4, 5.9 Hz, 1H), 4.43 (ddd, *J* = 10.1, 6.7, 3.1 Hz, 2H), 4.38 – 4.32 (m, 1H), 4.21 (dd, *J* = 15.9, 5.6 Hz, 1H), 3.71 – 3.61 (m, 2H), 3.25 (dd, *J* = 15.1, 8.3 Hz, 1H), 3.21 – 3.00 (m, 3H), 2.59 (s, 3H), 2.44 (s, 3H), 2.41 (s, 3H), 2.32 –2.24 (m, 1H), 2.21 – 2.09 (m, 1H), 2.08 – 1.98 (m, 1H), 1.90 (ddd, *J* = 12.9, 8.6, 4.6 Hz, 1H), 1.62 (s, 3H), 1.65 – 1.37 (m, 4H), 0.93 (s, 9H). ^13^C NMR (101 MHz, DMSO*-d6*) δ 171.9 (2C), 169.7, 169.3, 163.0, 155.1, 151.3 (visible via HSQC coupling), 149.8, 147.7, 139.5, 136.8, 135.2, 132.3, 131.1, 130.7, 130.1 (2C), 129.8, 129.6, 129.6, 128.6 (2C), 128.5 (2C), 127.4 (2C), 68.9, 58.7, 56.4, 56.3, 53.9, 41.6, 38.4, 37.9, 37.6, 35.2, 34.6, 28.9, 26.4 (3C), 23.0, 15.9, 14.1, 12.7, 11.3. HRMS, ESI+, m/z calcd for C_46_H_54_ClN_9_O_5_S_2_ [M+H]+, 912.3451; found 912.3447.

2-((6S)-4-(4-chlorophenyl)-2,3,9-trimethyl-6H-thieno[3,2-f][1,2,4]triazolo[4,3-a][1,4]diazepin-6-yl)-N-(((1RS,3SR)-3-(((2-(2,6-dioxopiperidin-3-yl)-1,3-dioxoisoindolin-4-yl)amino)methyl)cyclohexyl)methyl)acetamide (**12**).

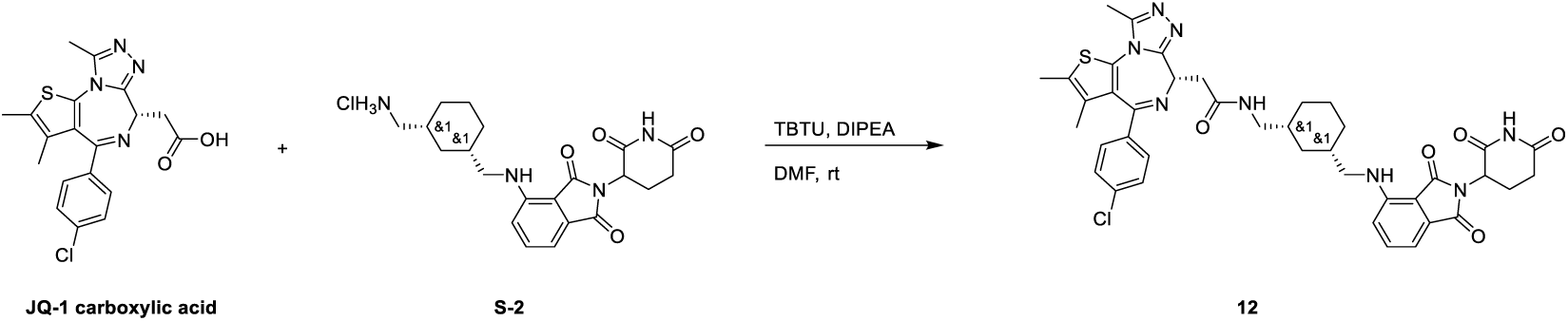

DIPEA (17 µL, 0.1 mmol) was added to a stirred solution of JQ-1 carboxylic acid (10 mg, 0.025 mmol), **S-2** (9.9 mg, 0.025 mmol) and TBTU (8.4 mg, 0.026 mmol) in DMF (0.5 mL). The reaction was allowed to perform at room temperature overnight. The mixture was diluted with EtOAc, washed with NaHCO_3_ sat. aq., dried over sodium sulfate, filtered and concentrated to dryness. The resulting material was purified using automated flash chromatography (12 g silica cartridge, DCM/MeOH: 0-10 % over 7 column volumes) to provide compound **12** (12.5 mg, 65 % yield). ^1^H NMR (400 MHz, DMSO*-d6*) δ 11.09 (s, 1H), 8.19 (q, *J* = 6.4 Hz, 1H), 7.56 (ddd, *J* = 9.4, 7.0, 2.5 Hz, 1H), 7.51 – 7.39 (m, 4H), 7.12 – 7.04 (m, 1H), 7.01 (d, *J* = 7.0 Hz, 1H), 6.63 – 6.52 (m, 1H), 5.05 (ddd, *J* = 12.9, 5.4, 1.8 Hz, 1H), 4.50 (dd, *J* = 8.4, 5.8 Hz, 1H), 3.35 – 3.10 (m, 4H), 3.08 – 2.80 (m, 3H), 2.64 – 2.50 (m, 5H), 2.41 (s, 3H), 2.08 – 1.97 (m, 1H), 1.85 – 1.69 (m, 4H), 1.66 – 1.55 (m, 4H), 1.55 – 1.40 (m, 1H), 1.28 – 1.14 (m, 1H), 0.97 – 0.78 (m, 2H), 0.66 (p, *J* = 12.7, 12.3 Hz, 1H). ^13^C NMR (101 MHz, DMSO*-d6*) δ 172.8, 170.1, 169.4, 169.0, 167.3, 163.0 (d), 155.1, 149.8, 146.6, 136.7, 136.2, 135.2, 132.3, 132.1, 130.7, 130.1 (d, 2C), 129.8, 129.6, 128.4 (2C), 117.3 (d), 110.4, 109.0, 54.0, 48.5, 48.2 (d), 44.9 (d), 37.7 (d), 37.6 (d), 37.4 (d), 37.0 (d), 34.6, 31.0, 30.4, 30.2 (d), 25.0, 22.2, 14.1 (d), 12.7, 11.3. HRMS, ESI+, m/z calcd for C_40_H_41_ClN_8_O_5_S [M+H]+, 781.2682; found 781.2677.

2-((6S)-4-(4-chlorophenyl)-2,3,9-trimethyl-6H-thieno[3,2-f][1,2,4]triazolo[4,3-a][1,4]diazepin-6-yl)-N-(3-((2-(2,6-dioxopiperidin-3-yl)-1,3-dioxoisoindolin-4-yl)amino)propyl)acetamide (**13**).

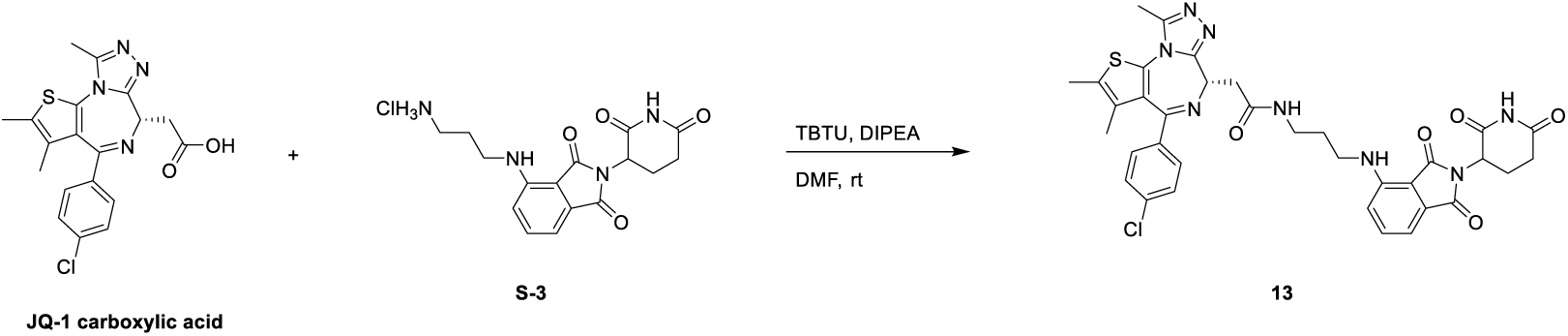

DIPEA (17 µL, 0.1 mmol) was added to a stirred solution of JQ-1 carboxylic acid (10 mg, 0.025 mmol), **S-3** (8.2 mg, 0.025 mmol) and TBTU (8.4 mg, 0.026 mmol) in DMF (0.5 mL). The reaction was allowed to perform at room temperature overnight. The mixture was diluted with EtOAc, washed with NaHCO_3_ sat. aq., dried over sodium sulfate, filtered and concentrated to dryness. The resulting material was purified using automated flash chromatography (12 g silica cartridge, DCM/MeOH: 0-10 % over 7 column volumes) to provide compound **13** (10.7 mg, 60 % yield). ^1^H NMR (400 MHz, DMSO*-d6*) δ 11.09 (s, 1H), 8.32 (t, *J* = 5.9 Hz, 1H), 7.56 (dd, *J* = 8.6, 7.0 Hz, 1H), 7.52 – 7.37 (m, 4H), 7.10 (d, *J* = 8.6 Hz, 1H), 7.02 (d, *J* = 7.0 Hz, 1H), 6.67 (td, *J* = 6.3, 2.1 Hz, 1H), 5.05 (dd, *J* = 12.8, 5.4 Hz, 1H), 4.52 (t, *J* = 7.2 Hz, 1H), 3.36 (q, *J* = 6.8 Hz, 2H), 3.29 – 3.16 (m, 4H), 2.88 (ddd, *J* = 17.8, 14.2, 5.7 Hz, 1H), 2.65 – 2.49 (m, 5H), 2.40 (s, 3H), 2.07 – 1.96 (m, 1H), 1.79 – 1.67 (m, 2H), 1.61 (s, 3H). ^13^C NMR (101 MHz, DMSO*-d6*) δ 172.8, 170.1, 169.7, 168.8, 167.3, 163.1, 155.1, 149.8, 146.3, 136.7, 136.2, 135.2, 132.3, 130.7, 130.1 (2C), 129.8, 129.6, 128.4 (2C), 117.2, 110.4, 109.2, 53.9, 48.5, 39.3, 37.8, 35.8, 35.8, 31.0, 28.9, 22.2, 14.0, 12.7, 11.3. HRMS, ESI+, m/z calcd for C_35_H_33_ClN_8_O_5_S [M+H]+, 713.2056; found 713.2052.

2-((6S)-4-(4-chlorophenyl)-2,3,9-trimethyl-6H-thieno[3,2-f][1,2,4]triazolo[4,3-a][1,4]diazepin-6-yl)-N-(2-(2-(2-(2-((2-(2,6-dioxopiperidin-3-yl)-1,3-dioxoisoindolin-4-yl)oxy)acetamido)ethoxy)ethoxy)ethyl)acetamide (**14**)

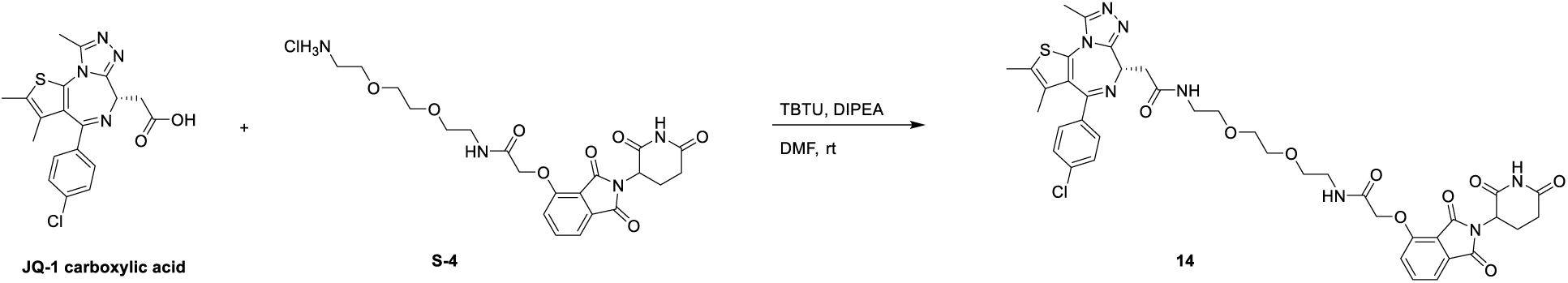

DIPEA (17 µL, 0.1 mmol) was added to a stirred solution of JQ-1 carboxylic acid (10 mg, 0.025 mmol), **S-4** (12 mg, 0.025 mmol) and TBTU (8.4 mg, 0.026 mmol) in DMF (0.5 mL). The reaction was allowed to perform at room temperature overnight. The mixture was diluted with EtOAc, washed with NaHCO_3_ sat. aq., dried over sodium sulfate, filtered and concentrated to dryness. The resulting material was purified using automated flash chromatography (12 g silica cartridge, DCM/MeOH: 0-10 % over 7 column volumes) to provide compound **14** (16 mg, 76 % yield). ^1^H NMR (400 MHz, DMSO*-d6*) δ 11.11 (s, 1H), 8.26 (t, *J* = 5.7 Hz, 1H), 8.02 (t, *J* = 5.7 Hz, 1H), 7.80 (dd, *J* = 8.5, 7.3 Hz, 1H), 7.51 – 7.36 (m, 6H), 5.11 (dd, *J* = 12.9, 5.5 Hz, 1H), 4.79 (s, 2H), 4.50 (dd, *J* = 8.0, 6.3 Hz, 1H), 3.58 – 3.52 (m, 4H), 3.46 (dt, *J* = 10.5, 5.9 Hz, 4H), 3.36 – 3.16 (m, 6H), 2.89 (ddd, *J* = 17.8, 14.3, 5.8 Hz, 1H), 2.64 – 2.53 (m, 5H), 2.40 (s, 3H), 2.09 – 1.98 (m, 1H), 1.61 (s, 3H). ^13^C NMR (101 MHz, DMSO*-d6*) δ 172.8, 169.9, 169.7, 166.9, 166.7, 165.4, 163.0, 155.1, 155.0, 149.8, 136.9, 136.8, 135.2, 133.0, 132.3, 130.7, 130.1 (2C), 129.8, 129.5, 128.4 (2C), 120.3, 116.8, 116.0, 69.6 (2C), 69.2, 68.8, 67.5, 53.8, 48.8, 38.6, 38.4, 37.5, 30.9, 22.0, 14.0, 12.7, 11.3. HRMS, ESI+, m/z calcd for C_40_H_41_ClN_8_O_9_S [M+H]+, 845.2479; found 845.2475.

4-(4-(2-((6S)-4-(4-chlorophenyl)-2,3,9-trimethyl-6H-thieno[3,2-f][1,2,4]triazolo[4,3-a][1,4]diazepin-6-yl)acetyl)piperazin-1-yl)-2-(2,6-dioxopiperidin-3-yl)isoindoline-1,3-dione (**15**).

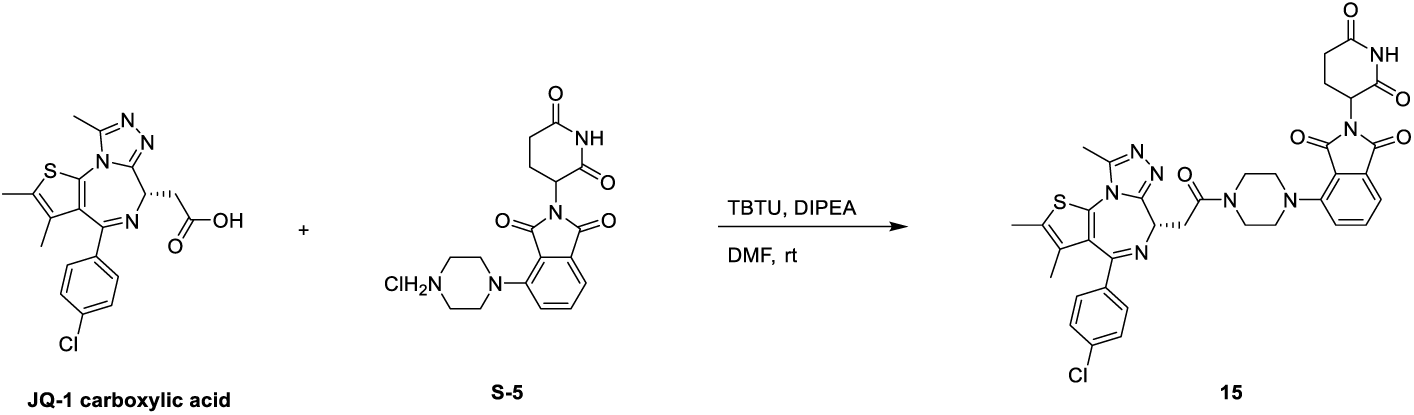

DIPEA (17 µL, 0.1 mmol) was added to a stirred solution of JQ-1 carboxylic acid (10 mg, 0.025 mmol), **S-5** (8.5 mg, 0.025 mmol) and TBTU (8.4 mg, 0.026 mmol) in DMF (0.5 mL). The reaction was allowed to perform at room temperature overnight. The mixture was diluted with EtOAc, washed with NaHCO_3_ sat. aq., dried over sodium sulfate, filtered and concentrated to dryness. The resulting material was purified using automated flash chromatography (12 g silica cartridge, DCM/MeOH: 0-10 % over 7 column volumes) to provide compound **15** (9.9 mg, 55 % yield). ^1^H NMR (400 MHz, DMSO-*d6*) δ 11.10 (s, 1H), 7.75 (dd, *J* = 8.4, 7.1 Hz, 1H), 7.55 – 7.32 (m, 6H), 5.13 (dd, *J* = 12.8, 5.4 Hz, 1H), 4.61 (t, *J* = 6.8 Hz, 1H), 3.96 – 3.81 (m, 2H), 3.77 – 3.61 (m, 3H), 3.52 – 3.37 (m, 3H), 3.32 – 3.23 (m, 2H), 2.89 (ddd, *J* = 17.5, 14.3, 5.6 Hz, 1H), 2.65 – 2.52 (m, 5H), 2.42 (s, 3H), 2.10 – 2.00 (m, 1H), 1.64 (s, 3H). ^13^C NMR (101 MHz, DMSO-*d6*) δ 172.8, 170.0, 168.5, 167.0, 166.4, 162.9, 155.2, 149.8, 149.4, 136.8, 136.0, 135.2, 133.6, 132.2, 130.7, 130.2 (2C), 129.9, 129.6, 128.5 (2C), 123.9, 117.0, 115.3, 54.2, 50.8, 50.4, 48.8, 45.0, 41.2, 34.8, 31.0, 22.0, 14.0, 12.7, 11.3. HRMS, ESI+, m/z calcd for C_36_H_33_ClN_8_O_5_S [M+H]+, 725.2056; found 725.2054.

2-((6S)-4-(4-chlorophenyl)-2,3,9-trimethyl-6H-thieno[3,2-f][1,2,4]triazolo[4,3-a][1,4]diazepin-6-yl)-N-(2-(2-(2-((2-(2,6-dioxopiperidin-3-yl)-1,3-dioxoisoindolin-4-yl)oxy)acetamido)ethoxy)ethyl)acetamide (**16**).

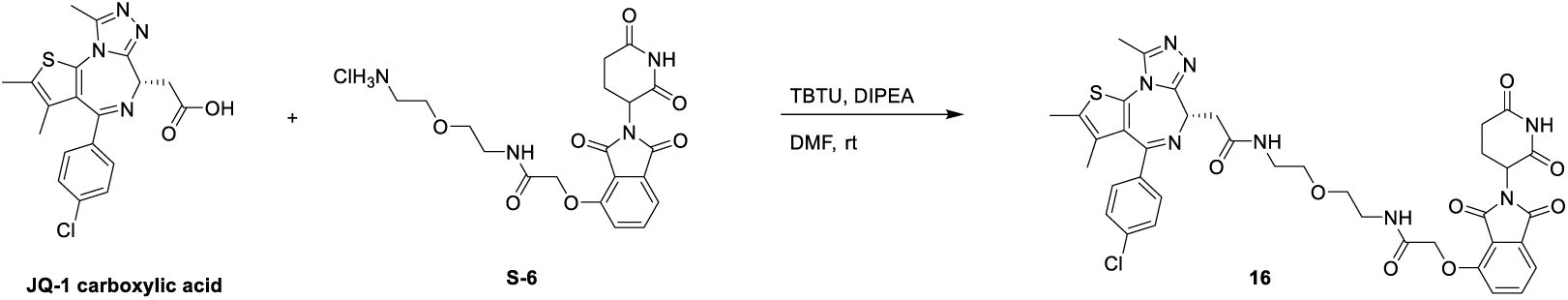

DIPEA (17 µL, 0.1 mmol) was added to a stirred solution of JQ-1 carboxylic acid (10 mg, 0.025 mmol), **S-6** (10 mg, 0.025 mmol) and TBTU (8.4 mg, 0.026 mmol) in DMF (0.5 mL). The reaction was allowed to perform at room temperature overnight. The mixture was diluted with EtOAc, washed with NaHCO_3_ sat. aq., dried over sodium sulfate, filtered and concentrated to dryness. The resulting material was purified using automated flash chromatography (12 g silica cartridge, DCM/MeOH: 0-10 % over 7 column volumes) to provide compound **16** (13 mg, 65 % yield). ^1^H NMR (400 MHz, DMSO-*d6*) δ 11.10 (s, 1H), 8.25 (t, *J* = 5.6 Hz, 1H), 8.04 (t, *J* = 5.6 Hz, 1H), 7.79 (dd, *J* = 8.6, 7.3 Hz, 1H), 7.52 – 7.33 (m, 6H), 5.11 (dd, *J* = 12.8, 5.3 Hz, 1H), 4.79 (s, 2H), 4.54 – 4.46 (m, 1H), 3.48 (dt, *J* = 11.3, 6.1 Hz, 4H), 3.38 – 3.17 (m, 6H), 2.88 (ddd, *J* = 17.6, 14.1, 5.6 Hz, 1H), 2.63 – 2.50 (m, 5H), 2.40 (s, 3H), 2.09 – 1.97 (m, 1H), 1.61 (s, 3H). ^13^C NMR (101 MHz, DMSO-*d6*) δ 172.8, 169.9, 169.7, 166.9, 166.7, 165.4, 163.0, 155.1, 155.0, 149.8, 136.9, 136.7, 135.2, 133.0, 132.2, 130.7, 130.1 (2C), 129.8, 129.5, 128.5 (2C), 120.3, 116.7, 116.0, 69.0, 68.6, 67.5, 53.8, 48.8, 38.5, 38.4, 37.5, 30.9, 22.0, 14.0, 12.7, 11.3. HRMS, ESI+, m/z calcd for C_38_H_37_ClN_8_O_8_S [M+H]+, 801.2217; found 801.2215.

4-((1-(2-((6S)-4-(4-chlorophenyl)-2,3,9-trimethyl-6H-thieno[3,2-f][1,2,4]triazolo[4,3-a][1,4]diazepin-6-yl)acetyl)pyrrolidin-3-yl)oxy)-2-(2,6-dioxopiperidin-3-yl)isoindoline-1,3-dione (**17**).

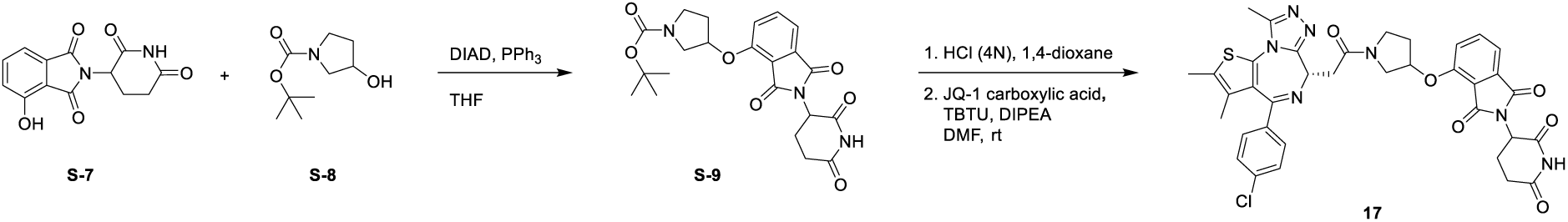

<u>tert-butyl 3-((2-(2,6-dioxopiperidin-3-yl)-1,3-dioxoisoindolin-4-yl)oxy)pyrrolidine-1-carboxylate (**S-9**).</u>

Tert-butyl 3-hydroxypyrrolidine-1-carboxylate (**S-8**) (162 mg, 0.87 mmol) and triphenylphosphine (526 mg, 2 mmol) were added to a stirred solution of 2-(2,6-dioxopiperidin-3-yl)-4-hydroxyisoindoline-1,3-dione (**S-7**) (250 mg, 0.91 mmol) in THF (10 mL). DIAD (358 µL) diluted in THF (6 mL) was then added dropwise to the mixture and the reaction was allowed to perform overnight at room temperature. The mixture was diluted with EtOAc, washed with water, dried over sodium sulfate, filtered and concentrated to dryness.

The resulting material was purified using automated flash chromatography (40 g silica cartridge, iso-Hexane/EtOAc, 1:3) to provide compound **S-9** (200 mg, 50 % yield). ^1^H NMR (400 MHz, MeOD-*d4*) δ 7.80 – 7.71 (m, 1H), 7.49 – 7.38 (m, 2H), 5.26 (dd, J = 3.9, 2.0 Hz, 1H), 5.09 (dd, J = 12.4, 5.6 Hz, 1H), 4.75 – 4.45 (m, 1H), 3.71 – 3.49 (m, 4H), 2.94 – 2.80 (m, 1H), 2.80 – 2.63 (m, 2H), 2.28 – 2.15 (m, 2H), 2.13 (ddd, J = 10.3, 5.4, 3.1 Hz, 1H), 1.49 – 1.39 (m, 9H). LC-MS, ESI+, m/z 344.1 [M+H-Boc]+.

<u>4-((1-(2-((6S)-4-(4-chlorophenyl)-2,3,9-trimethyl-6H-thieno[3,2-f][1,2,4]triazolo[4,3-a][1,4]diazepin-6-yl)acetyl)pyrrolidin-3-yl)oxy)-2-(2,6-dioxopiperidin-3-yl)isoindoline-1,3-dione (**17**).</u>

Step 1: HCl (4N, in 1,4-dioxane) (6 mL, 24 mmol) was added to a stirred solution of **S-9** (200 mg, 0.45 mmol) in 1,4-dioxane (4 mL). After 3h, the heterogeneous solution was diluted with iso-hexane and filtered through a P3 glass-silica pad. The white solid was dried under vacuum to provide 2-(2,6-dioxopiperidin-3-yl)-4-(pyrrolidin-3-yloxy)isoindoline-1,3-dione HCl salt (140 mg, 94% yield).

Step 2: DIPEA (17 µL, 0.1 mmol) was added to a stirred solution of JQ-1 carboxylic acid (10 mg, 0.025 mmol), 2-(2,6-dioxopiperidin-3-yl)-4-(pyrrolidin-3-yloxy)isoindoline-1,3-dione (8.6 mg, 0.025 mmol) and TBTU (8.4 mg, 0.026 mmol) in DMF (0.5 mL). The reaction was allowed to perform at room temperature overnight. The mixture was diluted with EtOAc, washed with NaHCO_3_ sat. aq., dried over sodium sulfate, filtered and concentrated to dryness. The resulting material was purified using automated flash chromatography (12 g silica cartridge, DCM/MeOH: 0-10 % over 7 column volumes) to provide compound **17** (13.5 mg, 75 % yield) as a mixture of diastereoisomers (2/3 ratio).

Major diastereoisomer: ^1^H NMR (400 MHz, DMSO-*d6*) δ 11.12 (s, 1H), 7.84 (d, *J* = 7.3 Hz, 1H), 7.60 (d, *J* = 8.6 Hz, 1H), 7.56 – 7.32 (m, 5H), 5.39 – 5.30 (m, 1H), 5.14 – 5.04 (m, 1H), 4.61 – 4.52 (m, 1H), 4.01 (td, *J* = 9.9, 9.0, 2.7 Hz, 1H), 3.87 – 3.77 (m, 1H), 3.75 – 3.56 (m, 2H), 3.53 – 3.33 (m, 2H), 2.96 – 2.81 (m, 1H), 2.65 – 2.51 (m, 5H), 2.42 (s, 3H), 2.40 – 2.12 (m, 2H), 2.10 – 1.97 (m, 1H), 1.64 (s, 3H). ^13^C NMR (101 MHz, DMSO-*d6*) δ 173.2, 170.4, 168.8, 167.2, 165.7, 163.5, 155.7, 154.8, 150.3, 137.5, 137.3, 135.6, 134.0, 132.7, 131.2, 130.7 (2C), 130.4, 130.1, 128.9 (2C), 122.0, 117.9, 116.4, 77.5, 54.5, 52.1, 49.2, 44.8, 36.8, 31.8, 31.4, 22.5, 14.5, 13.2, 11.8.

Minor diastereoisomer: ^1^H NMR (400 MHz, DMSO-*d6*) δ 11.10 (s, 1H), 7.85 (d, *J* = 7.3 Hz, 1H), 7.64 (d, *J* = 8.6 Hz, 1H), 7.56 – 7.32 (m, 5H), 5.47 – 5.40 (m, 1H), 5.14 – 5.04 (m, 1H), 4.61 – 4.52 (m, 1H), 4.13 (dd, *J* = 11.9, 4.5 Hz, 1H), 3.95 – 3.87 (m, 1H), 3.75 – 3.56 (m, 2H), 3.53 – 3.33 (m, 2H), 2.96 – 2.81 (m, 1H), 2.65 – 2.51 (m, 5H), 2.42 (s, 3H), 2.40 – 2.12 (m, 2H), 2.10 – 1.97 (m, 1H), 1.64 (s, 3H). ^13^C NMR (101 MHz, DMSO-*d6*) δ 173.2, 170.4, 168.6, 167.2, 165.7, 163.4, 155.6, 154.8, 150.3, 137.4, 137.2, 135.6, 134.0, 132.6, 131.2, 130.6 (2C), 130.3, 130.1, 129.0 (2C), 121.9, 117.8, 116.4, 78.9, 54.3, 51.6, 49.2, 44.0, 36.9, 31.8, 30.5, 22.5, 14.5, 13.2, 11.8.

HRMS, ESI+, m/z calcd for C_36_H_32_ClN_7_O_6_S [M+H]+, 726.1896; found 726.1890.

### Statistics

Statistical analyses were performed in R (v.4.2.2). The goodness of fit for the linear regressions were determined by calculating the coefficient of determination (R^2^) with the R package ggpmisc (v 0.6.0).

## Acknowledgments

This work was supported by the Swedish Research Council, grant number 2018-03288 and Magn. Bergvalls Stiftelse, grant number 2020-04065.

## Author Contributions

E.I performed *in vitro* and *in silico* experiments, analyzed the data and wrote the paper. M.E provided technical assistance with *in vitro* experiments. L.A provided technical assistance with mass spectrometry analysis. H.L provided technical assistance with molecular dynamics simulations. R.C and F.K performed chemical synthesis and compound characterization. C.H. contributed to study conceptualization and reviewed and edited the paper. P.M conceptualized and initiated the study, provided funding and reviewed and edited the paper.

## Data availability

Supplemental data are available in the Zenodo repository, as part of record https://doi.org/10.5281/zenodo.14629078, and are divided into five files: ’Molecular formula strings.xlsx’ contains SMILES format structures and separation into E3 ligase ligand, linker and POI ligand substructures for all 3589 analyzed PROTACs, and their original literature references; ’2D Key experimental details.xlsx’ contains molecular properties calculated from 2D structures (used in Table 1); ’3D Key experimental details.xlsx’ contains molecular properties calculated from 3D structures (Figure 2); ’Membrane simulation details.xlsx’ contains details on the PROTAC–membrane simulations (Figures 3-5); and, ’Experimental key details.xlsx’ contains details on the experimental measurements of PROTAC–cell membrane interactions (Figures 3 and 5). All other relevant data generated and analyzed during this study are included in this article and its supplementary information.

