## Supplemental material for "Conformational dynamics in the membrane interactions of bispecific targeted degrader therapeutics"

Table of Contents

**Supplementary Tables**

Supplementary Table 1 ........ S3
Supplementary Table 2 ........ S3
Supplementary Table 3 ........ S4
Supplementary Table 4 ........ S4

**Supplementary Figures**

Supplementary Figure 1 ........ S5
Supplementary Figure 2 ........ S6
Supplementary Figure 3 ........ S6
Supplementary Figure 4 ........ S7
Supplementary Figure 5 ........ S7
Supplementary Figure 6 ........ S8
Supplementary Figure 7 ........ S9

**NMR spectra**

1H-, 13C- and HSQC-NMR spectra of compound **11** ........ S10
1H- and 13C-NMR spectra of compound **12** ........ S12
1H- and 13C-NMR spectra of compound **13** ........ S13
1H- and 13C-NMR spectra of compound **14** ........ S14
1H- and 13C-NMR spectra of compound **15** ........ S15
1H- and 13C-NMR spectra of compound **16** ........ S16
1H-NMR spectra of compound **S-9** ........ S17
1H- and 13C-NMR spectra of compound **17** ........ S18

#### Supplementary Tables

**Supplementary Table 1**. Summary of molecular descriptors (median (min-max range)) of full PROTAC molecules, and for the separated E3 ligand, linker and POI ligand domains for PROTACs reported to bind to other E3 ligases than CRBN and VHL.

| **PROTACs binding to other E3 ligases** | | | | |
|---|---|---|---|---|
| | **Full molecule** | **E3 ligand** | **Linker** | **POI ligand** |
| **n types** | 77 | 32 | 59 | 35 |
| **MW[a] [Da]** | 1051 (774-1514) | 482 (271-664) | 190 (44-440) | 394 (281-860) |
| **cLogP[b]** | 7.5 (2.1-14) | 2.5 (0.5-7.5) | 0 (-1.3-3.9) | 3.8 (1.3-8.6) |
| **TPSA[c] [Å$^2$]** | 209 (130-281) | 92 (26-138) | 45 (0-103) | 68 (42-116) |
| **HBD[d]** | 5 (1-7) | 3 (0-4) | 0 (0-2) | 1 (0-2) |
| **HBA[e]** | 14 (9-20) | 5 (2-9) | 4 (0-10) | 6 (3-9) |
| **nRotB[f]** | 25 (14-36) | 8 (1-11) | 10 (1-21) | 5 (1-13) |

[a] molecular weight; [b] calculated LogP; [c] topological surface area; [d] hydrogen bond donors;
[e] hydrogen bond acceptors; [f] number of rotatable bonds

**Supplementary Table 2**. Summary statistics of conformation-dependent molecular properties (n = 891 PROTACs). Values are presented as medians (min-max ranges).

| | **All PROTACs** | | |
|---|---|---|---|
| **n conformations in chloroform** | 25904 | | |
| **n conformations in water** | 43438 | | |
| | **CRBN[c]-binding** | **VHL[d]-binding** | **Other** |
| **n PROTACs** | 477 | 391 | 23 |
| **TPSA$_{\text{full molecule}}$[a]** | 219 (96-793) Å$^2$ | 219 (124-353) Å$^2$ | 225 (183-257) Å$^2$ |
| **TPSA$_{\text{E3 ligand}}$[a]** | 96 (58-175) Å$^2$ | 112 (95-152) Å$^2$ | 103 (62-120) Å$^2$ |
| **3D SA PSA[b] chloroform** | 203 (50-536) Å$^2$ | 149 (41-319) Å$^2$ | 140 (65-233) Å$^2$ |
| **3D SA PSA[b] water** | 256 (76-716) Å$^2$ | 193 (44-469) Å$^2$ | 180 (81-298) Å$^2$ |
| **% 3D SA PSA[b] of TSA[c] in chloroform** | 20% (5%-44%) | 13% (4%-28%) | 12% (6%-18%) |
| **% 3D SA PSA[b] of TSA[c] in water** | 24% (7%-50%) | 17% (6%-38%) | 15% (10%-24%) |
| **Difference water-chloroform** | 50 (-64-237) Å$^2$ | 41 (-22-196) Å$^2$ | 35 (-21-114) Å$^2$ |
| **Maximum shielded 3D SA PSA[b]** | 116 (6-300) Å$^2$ | 97 (25-248) Å$^2$ | 92 (48-190) Å$^2$ |
| **Normalized shielded 3D SA PSA[b]** | 40% (2%-80%) | 43% (14%-76%) | 42% (24%-70%) |

[a] topological surface area; [b] solvent accessible polar surface area; [c] solvent accessible total surface area;
[c] cereblon; [d] von Hippel-Lindau

**Supplementary Table 3**. Summary statistics of the ability of PROTAC molecules (n = 157) to hide polar surface area. The difference in PSA from membrane core to water has been calculated from the Boltzmann weighted conformer ensembles.

| **Group** | **% of molecules** | **Difference in PSA from membrane core to water** |
|---|---|---|
| PSA and TSA higher in membrane core than in water | 40.8% | 0-82 Å$^2$ |
| PSA higher and TSA lower in membrane core than in water | 4.9% | 0-7 Å |
| PSA lower and TSA higher in membrane core than in water | 27.2% | -36-0 Å$^2$ |
| PSA and TSA lower in membrane core than in water | 27.2% | -102-0 Å$^2$ |
| > 10 Å$^2$ higher PSA in membrane core than in water | 10.9% | |
| 0-10 Å$^2$ higher PSA in membrane core than in water | 34.7% | |
| 0-10 Å$^2$ lower PSA in membrane core than in water | 38.1% | |
| > 10 Å$^2$ lower PSA in membrane core than in water | 16.2% | |

**Supplementary Table 4**. Mass transitions, cone voltages, collision energies.

| **Compound** | **ESI mode** | **Charge** | **Parent** | **Daughter** | **Cone (V)** | **Collision (eV)** |
|---|---|---|---|---|---|---|
| 1 | ESI+ | +2 | 560.9951 | 318.2010 | 34 | 16 |
| 2 | ESI+ | +2 | 512.9183 | 318.0852 | 16 | 28 |
| 3 | ESI+ | +2 | 499.7578 | 127.1340 | 22 | 22 |
| 4 | ESI+ | +2 | 557.4903 | 318.0979 | 16 | 16 |
| 5 | ESI+ | +2 | 469.4172 | 358.0882 | 16 | 16 |
| 6 | ESI+ | +2 | 501.9786 | 188.1866 | 16 | 28 |
| 7 | ESI+ | +2 | 390.8361 | 309.1414 | 16 | 22 |
| 8 | ESI+ | +1 | 841.2766 | 383.0578 | 10 | 46 |
| 9 | ESI+ | +1 | 785.2534 | 383.0003 | 10 | 40 |
| 10 | ESI+ | +2 | 392.8116 | 309.1384 | 28 | 22 |
| 11 | ESI+ | +2 | 457.1584 | 318.1621 | 22 | 16 |
| 12 | ESI+ | +1 | 781.4137 | 383.1128 | 10 | 34 |
| 13 | ESI+ | +1 | 713.3591 | 383.0981 | 22 | 34 |
| 14 | ESI+ | +1 | 845.3931 | 383.1785 | 16 | 40 |
| 15 | ESI+ | +1 | 725.3309 | 383.1176 | 22 | 34 |
| 16 | ESI+ | +1 | 801.3659 | 341.1367 | 28 | 46 |
| 17 | ESI+ | +1 | 726.3359 | 383.1140 | 16 | 34 |
| Lopinavir | ESI+ | +1 | 629.4825 | 155.1768 | 40 | 46 |
| Atorvastatin | ESI+ | +1 | 559.3015 | 440.1533 | 34 | 22 |
| Verapamil | ESI+ | +1 | 455.2701 | 165.0336 | 70 | 28 |

### Supplementary Figures

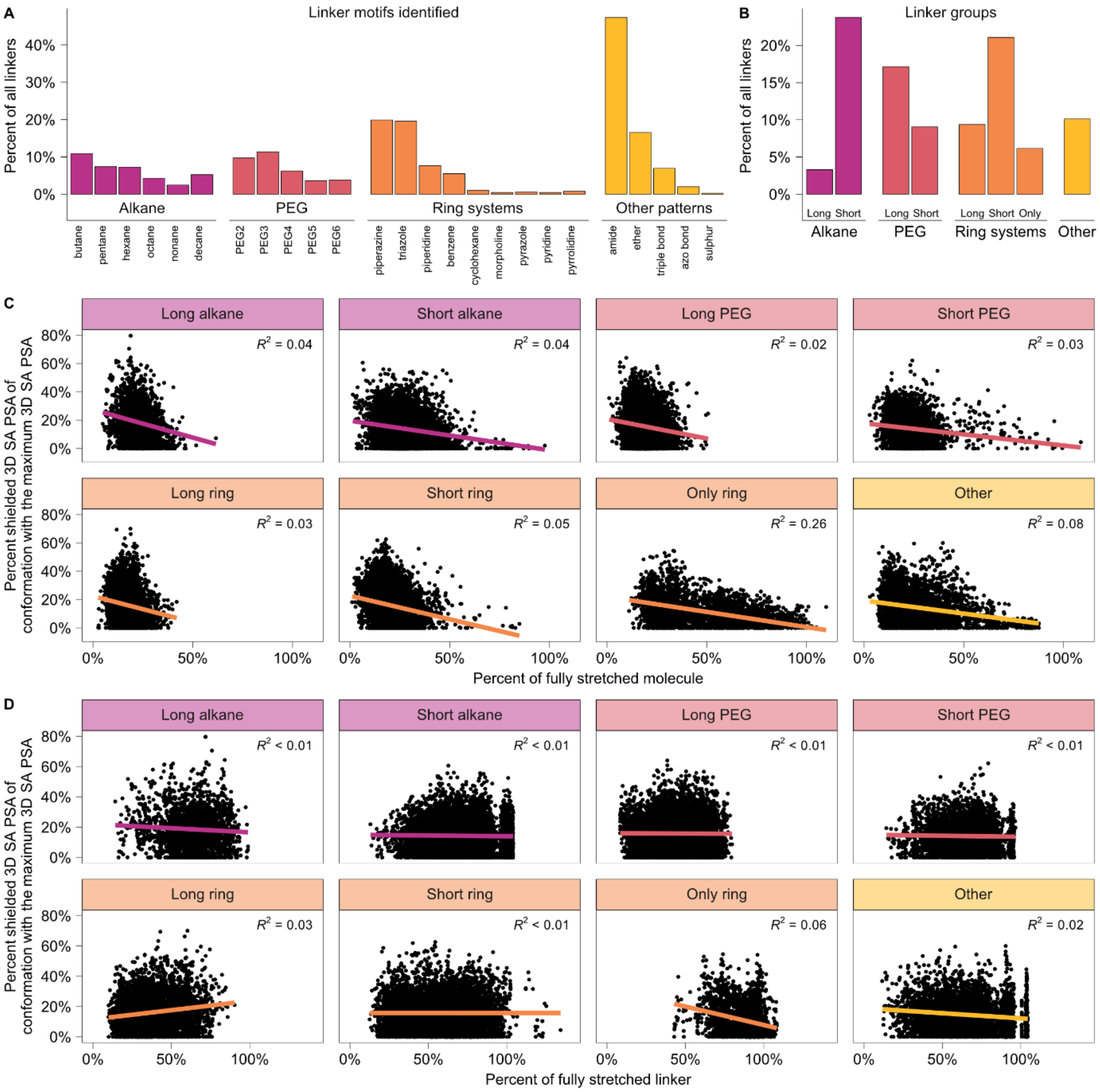

**Supplementary Figure 1**. Linker characteristics and origins and impact of PROTAC folding. A) Linker patterns identified in the dataset (n = 3576). One linker fragment can contain several linker patterns, or none, leading to cumulative percentages that can exceed 100%. B) Linker groups based on chain length and chemistry; long/short alkane (≥9/<9 carbons), long/short PEG (≥3/<3 PEG monomers), long/short ring systems (any linker containing a ring in combination with a long/short alkane or PEG segment), only ring systems (no alkane or PEG segment attached to ring) and other (linkers that fit to several groups or to none). Each linker (n = 3576) was only assigned to a single group. C) Percent shielding of each type of linker group indicates that more shielding occurs in molecules which are folded to a higher degree (n = 891). D) No correlation seen between shielding of PSA and linker folding (n = 891).

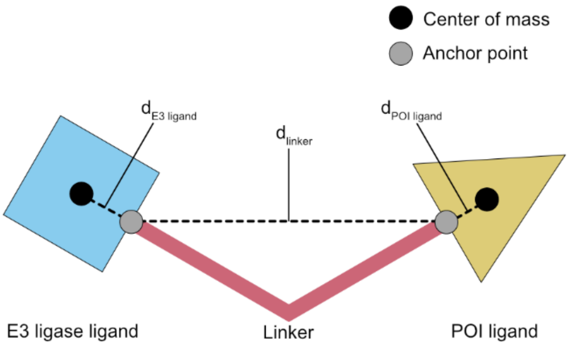

**Supplementary Figure 2**. Graphical illustration of how the linker and molecule contraction term were calculated. The sum of distances *E3 ligase center of mass to E3 ligase ligand anchor point* ($d_{E3\ ligand}$), *E3 ligase ligand anchor point to POI ligand anchor point* ($d_{linker}$) and *POI ligand anchor point to POI ligand center of mass* ($d_{POI\ ligand}$) was calculated for 2D and 3D conformations. Molecule contraction was calculated using Equation 2 and linker contraction was calculated using Equation 3 (see Methods section in the main article).

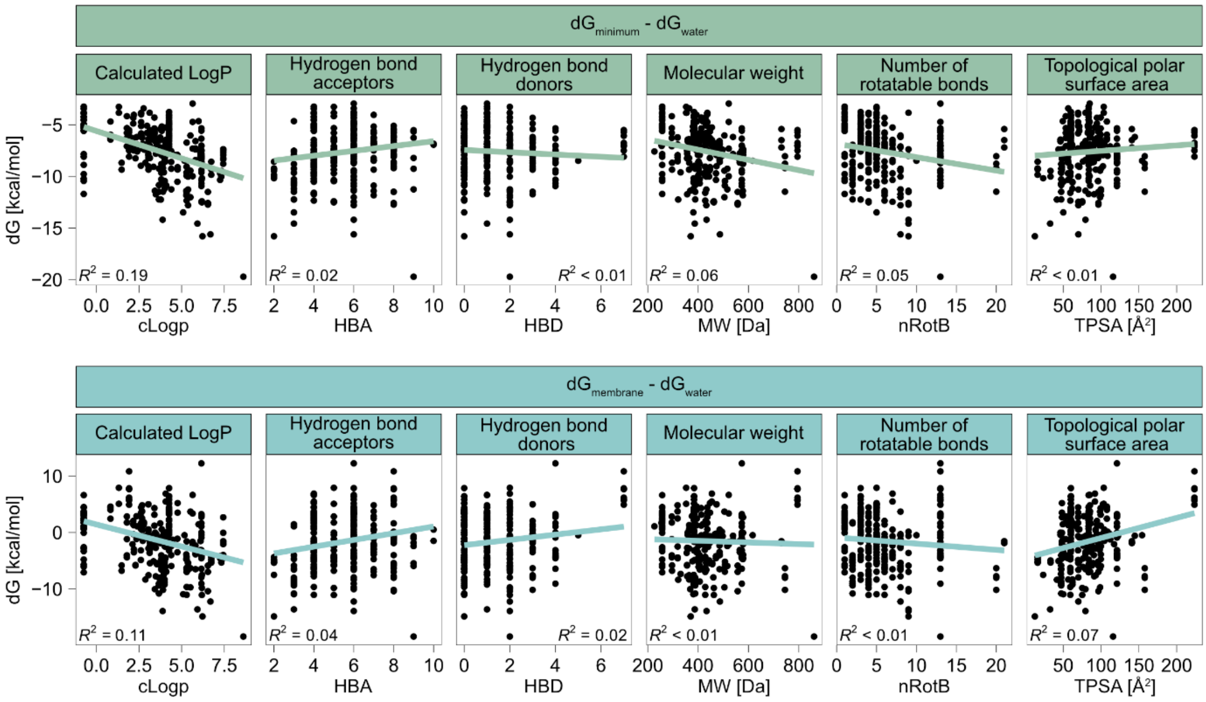

**Supplementary Figure 3**. Correlations between POI ligand characteristics and energy barriers (n =157).

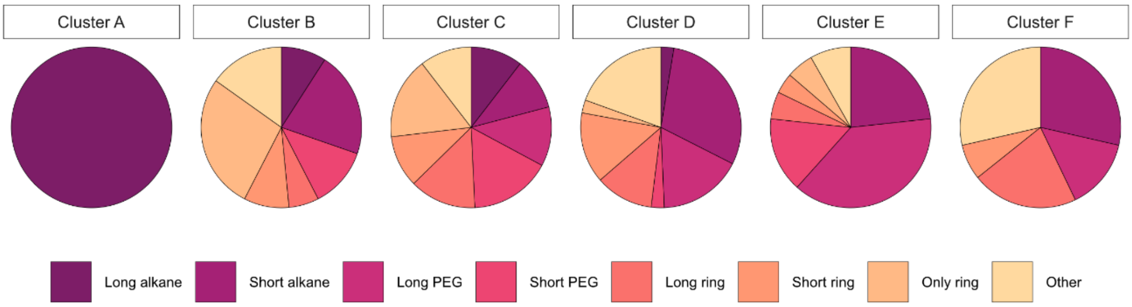

**Supplementary Figure 4**. Cluster characteristics (n = 157 PROTACs). Long PEG-based linkers are more common in clusters D-F, which have a higher energy barrier at the bilayer center. In contrast, more rigid and alkane-based linkers are more common in clusters A-C.

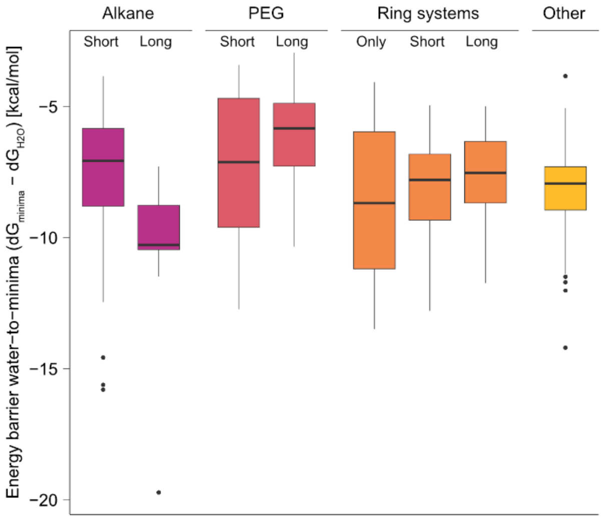

**Supplementary Figure 5**. Depth of energy minima as a function of linker chemistry (n = 157 PROTACs). On average, linkers containing long alkanes (≥9 carbons) result in deeper energy minima, as do linkers consisting solely of ring systems. Black solid line indicates the median and the bottom and upper box limits are the 25th and 75th percentile. Whiskers extend to 1.5 multiples of the interquartile length (IQR) and dots represent observations that extend outside of 1.5 IQR.

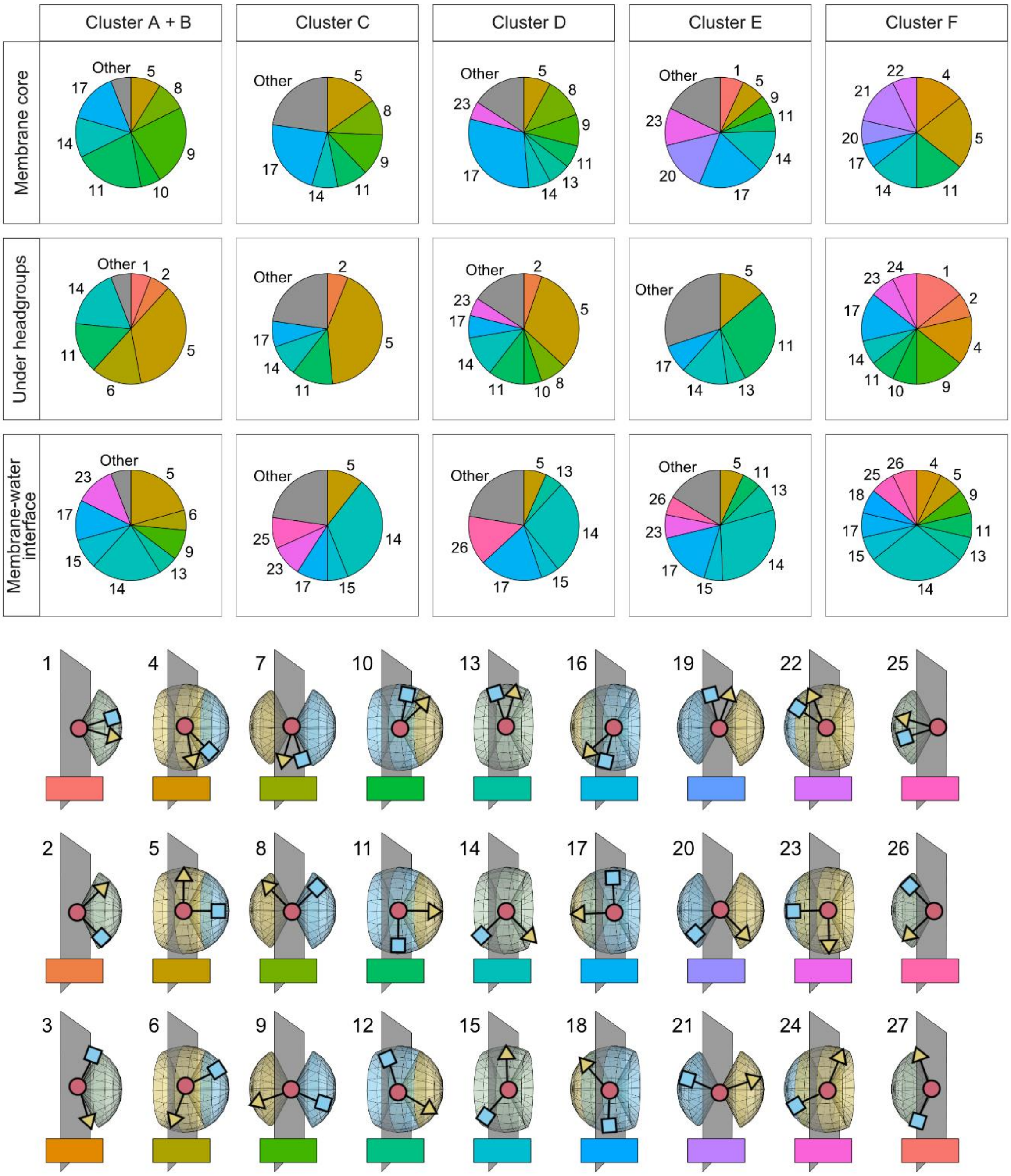

**Supplementary Figure 6**. PROTAC angles in the different clusters (n = 157 PROTACs). Clusters A and B were combined to one group. All orientation groups that more than 5% of molecules in each cluster belong to are indicated by a number, and those which contain less than 5% of molecules in each cluster are grouped together and marked Other. In clusters with deeper energy minima (cluster A + B), a preference for extended, more linear conformations are seen, with the POI ligand pointing in towards the core and ligase pointing towards the water. In clusters with higher energy barriers at the bilayer center extended conformations dominate, but with the ligase pointing in towards the core.

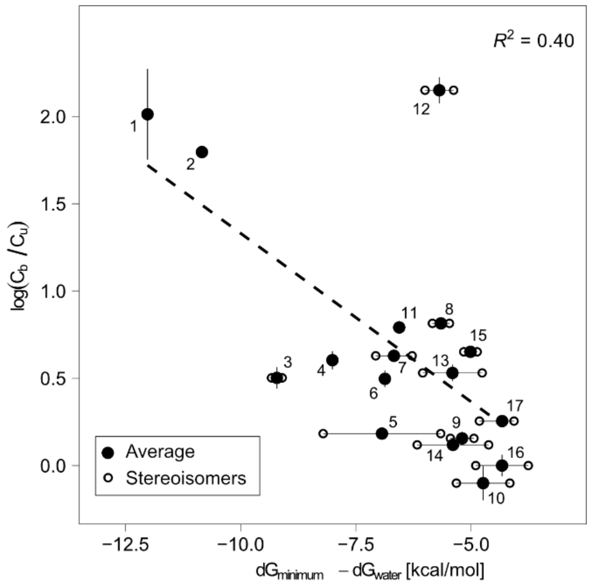

**Supplementary Figure 7**. Correlation between energy minimum and experimental membrane binding, expressed as the logarithm of bound-to-unbound concentration ratios (mean ± SD, n = 3). Full circles show the average of a 1:1 racemic mixture and open circles show values for the individual stereoisomers.

#### NMR spectra

1H-, 13C- and HSQC-NMR spectra of compound **11.**

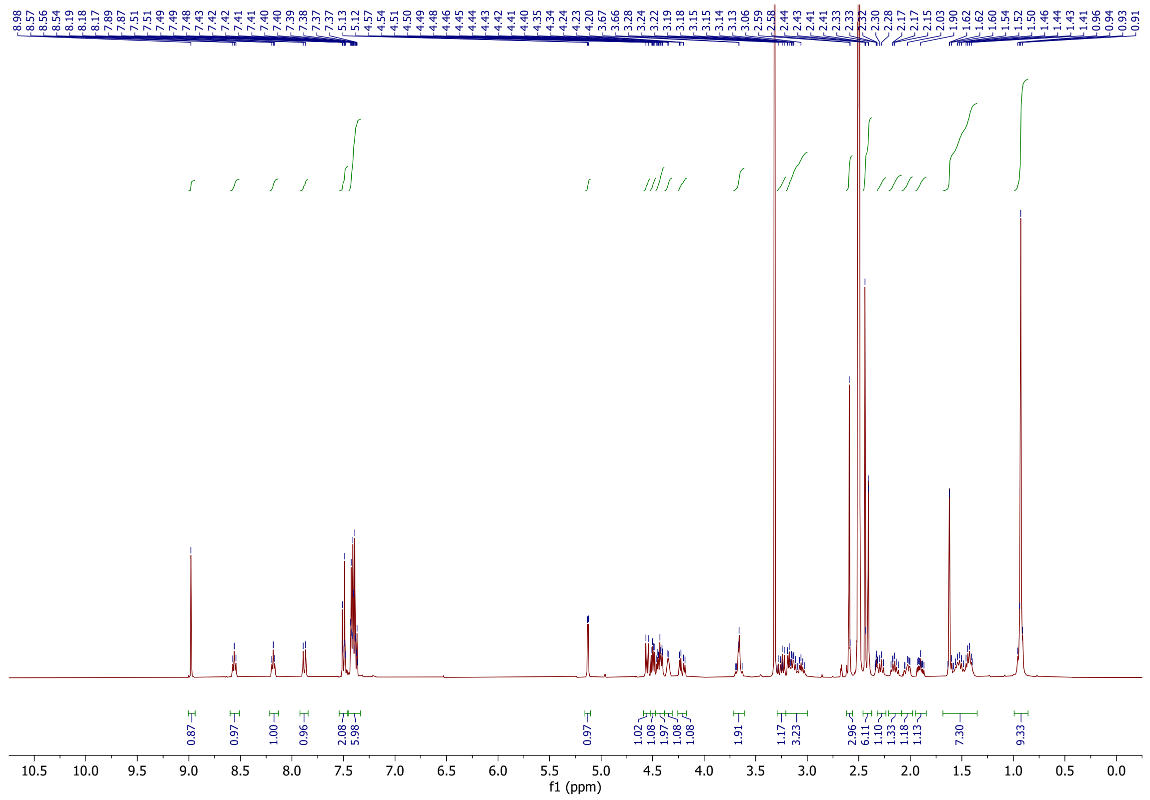

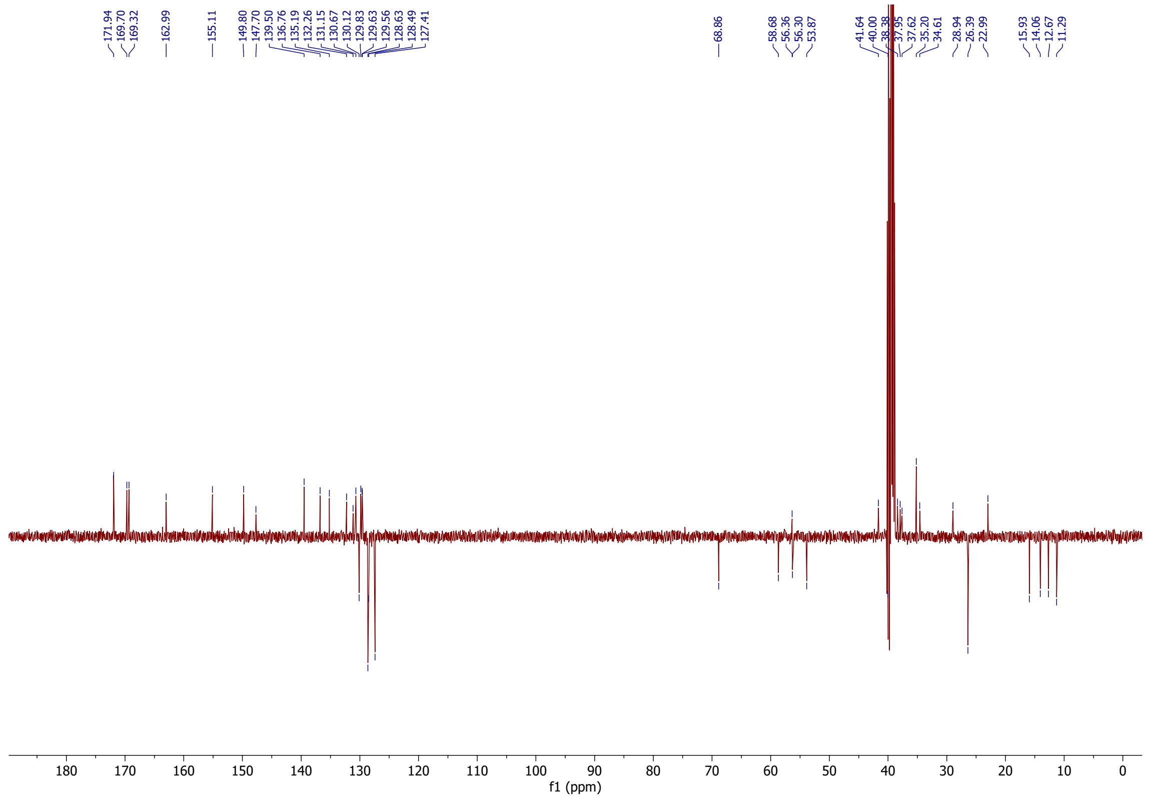

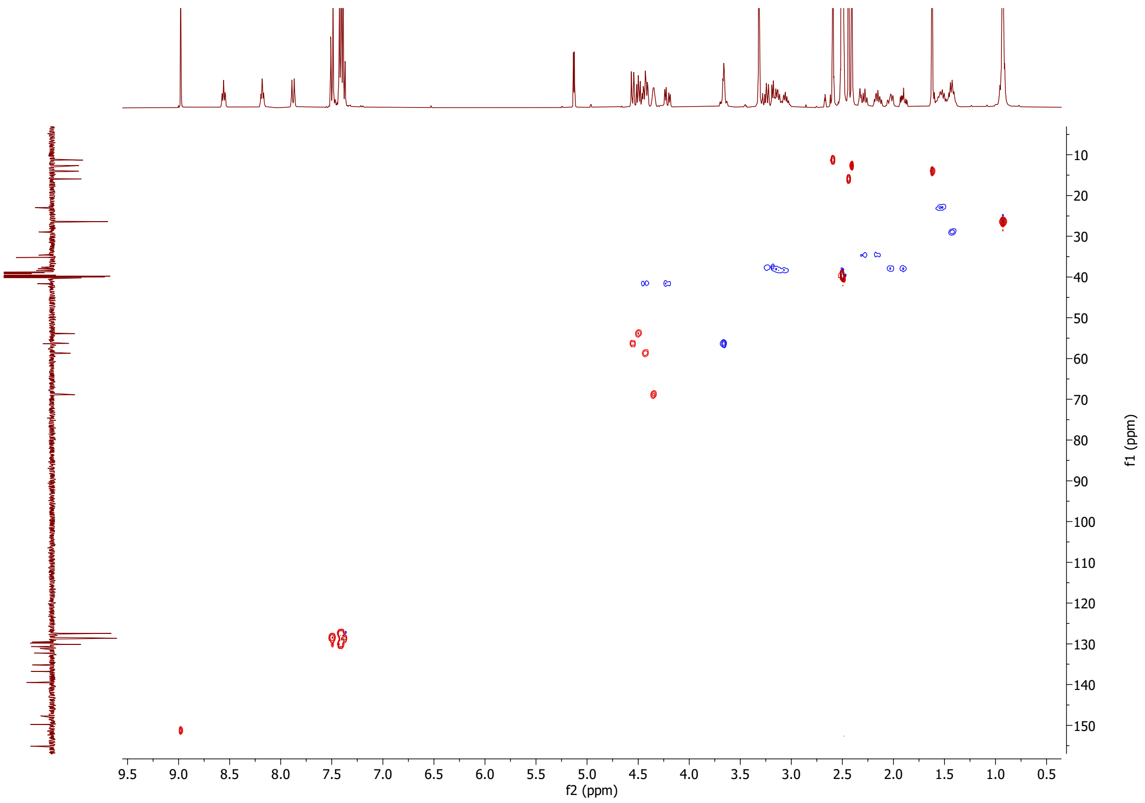
f1 (ppm)
f2 (ppm)

1H- and 13C-NMR spectra of compound **12**

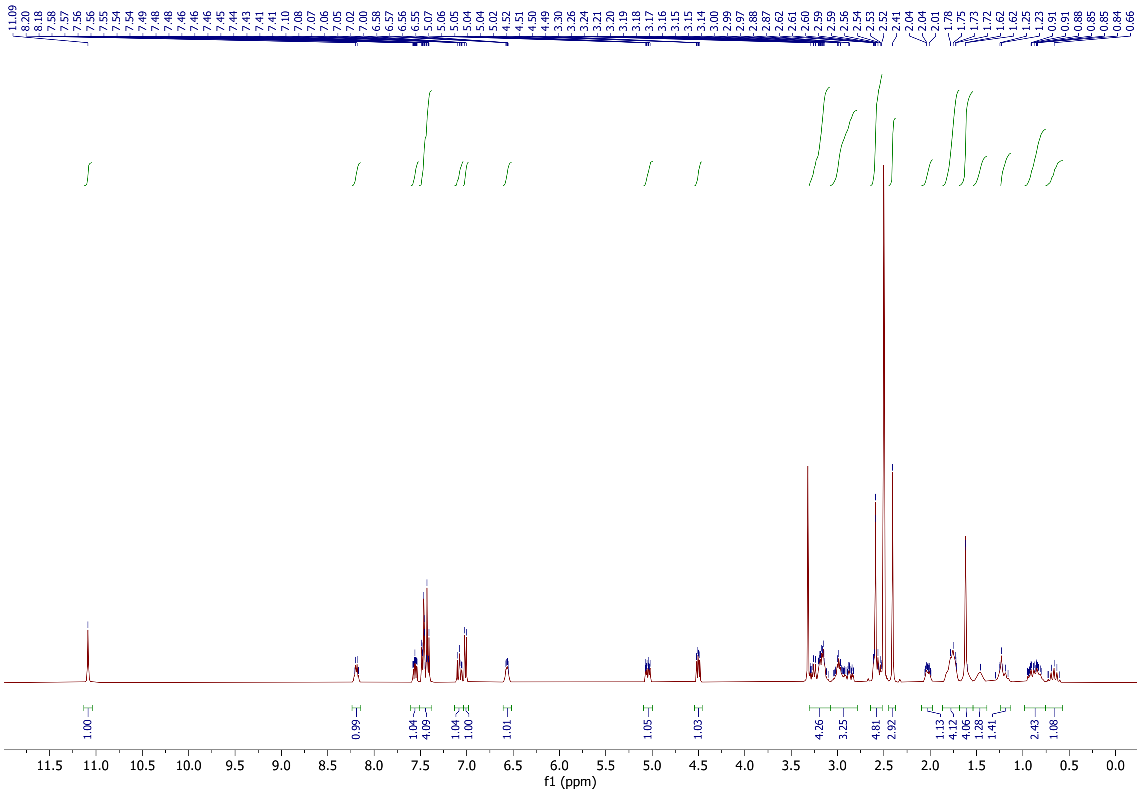

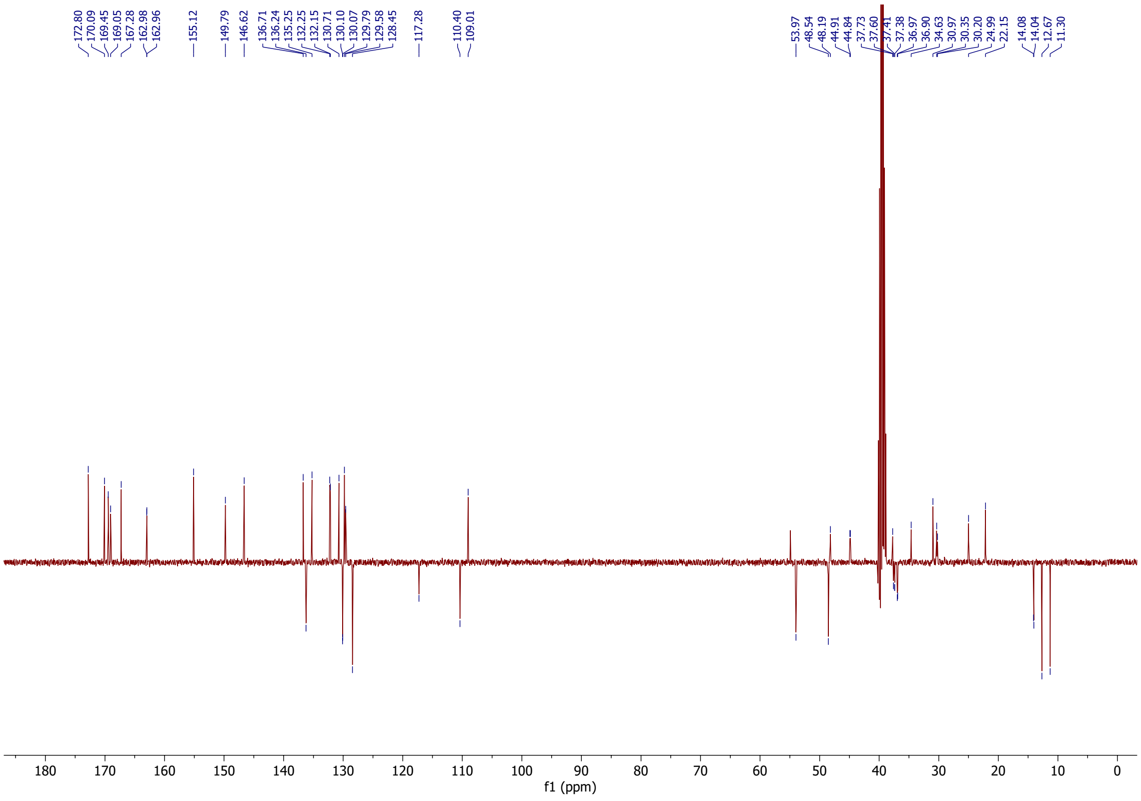

1H- and 13C-NMR spectra of compound **13**

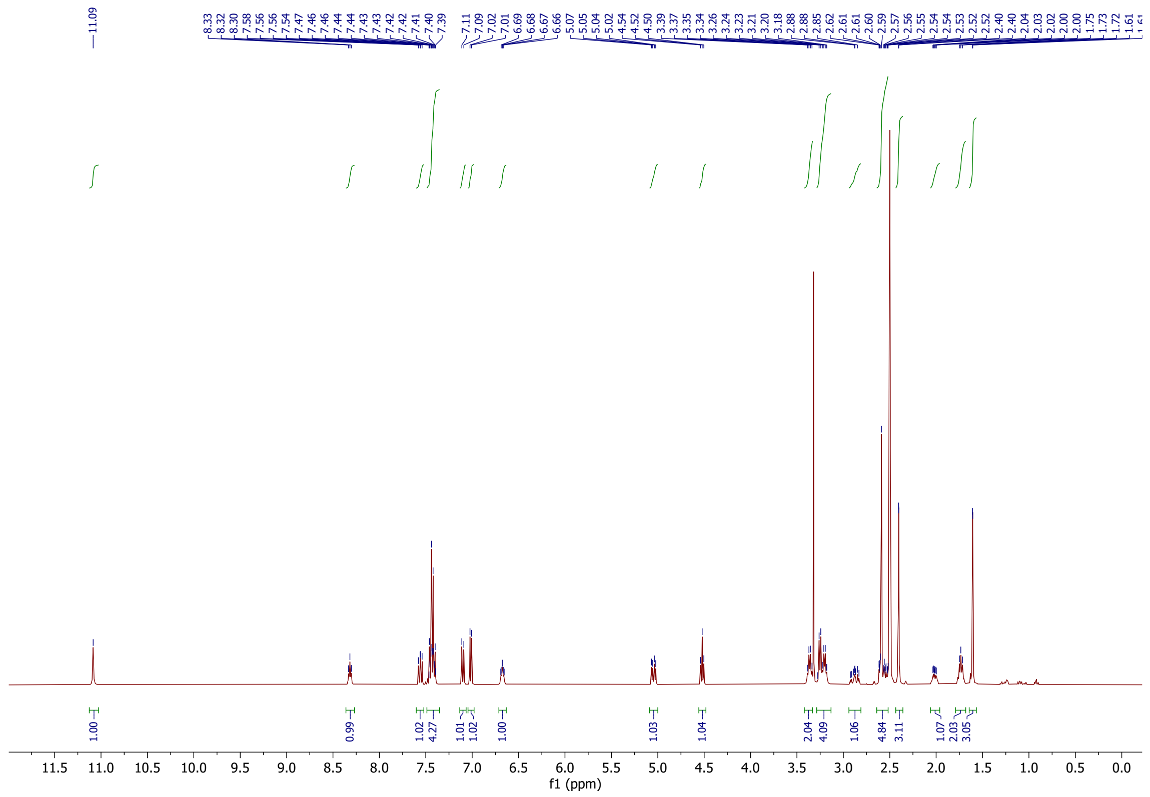

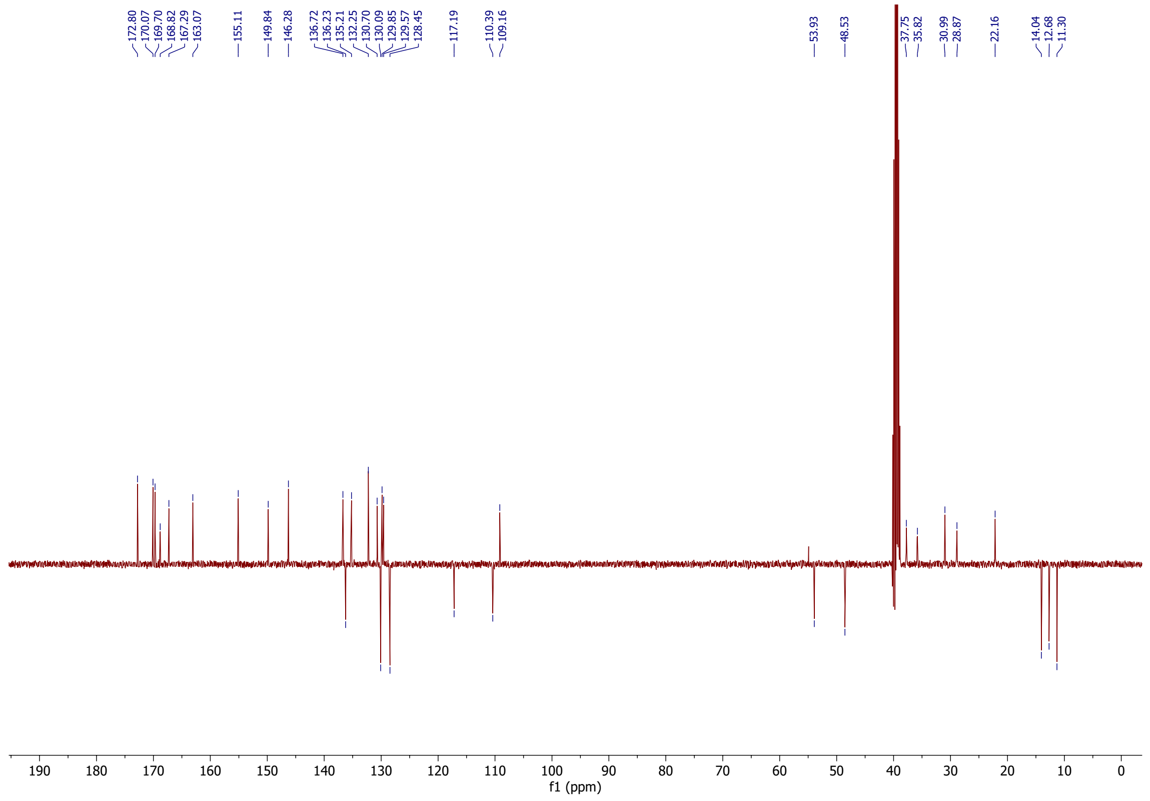

1H- and 13C-NMR spectra of compound **14**

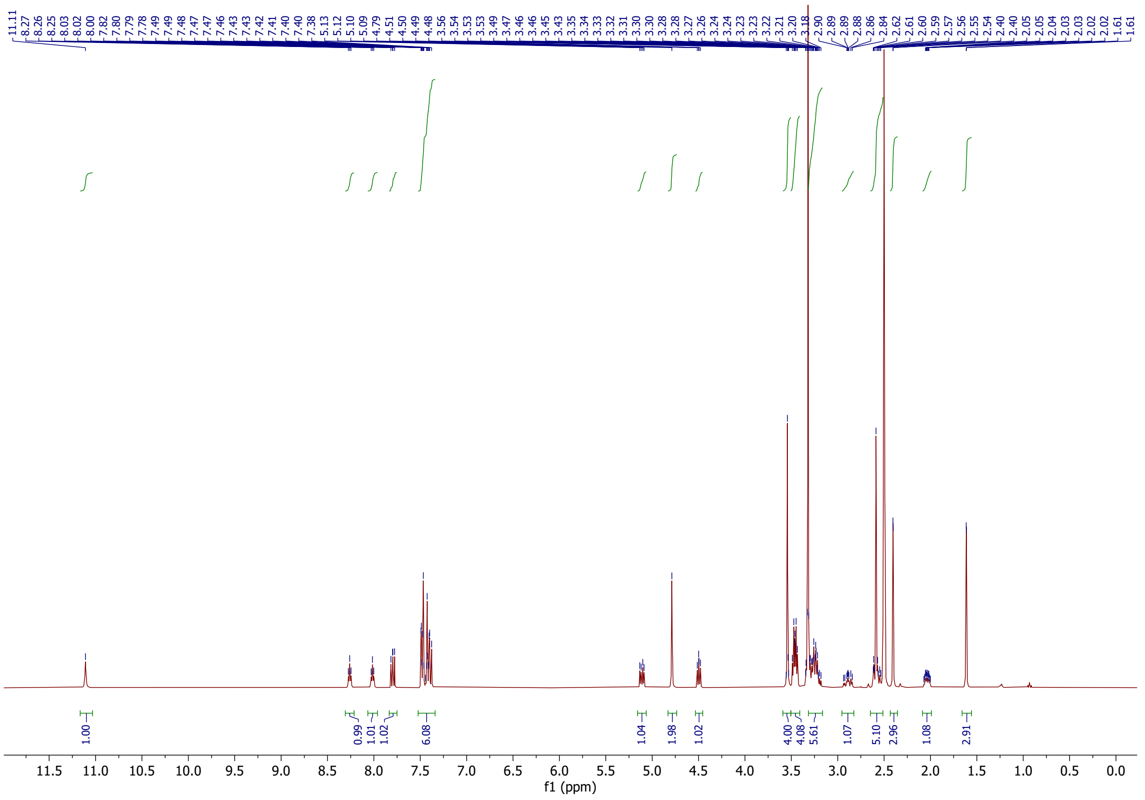

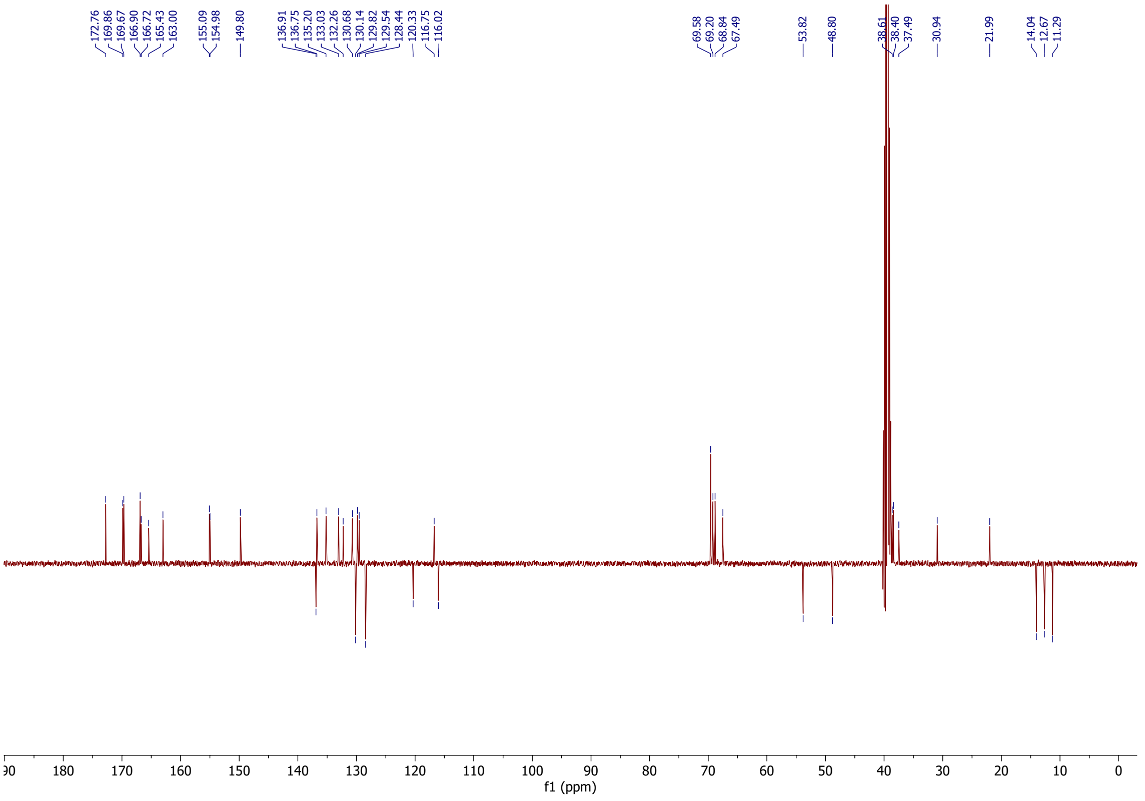

1H- and 13C-NMR spectra of compound **15**

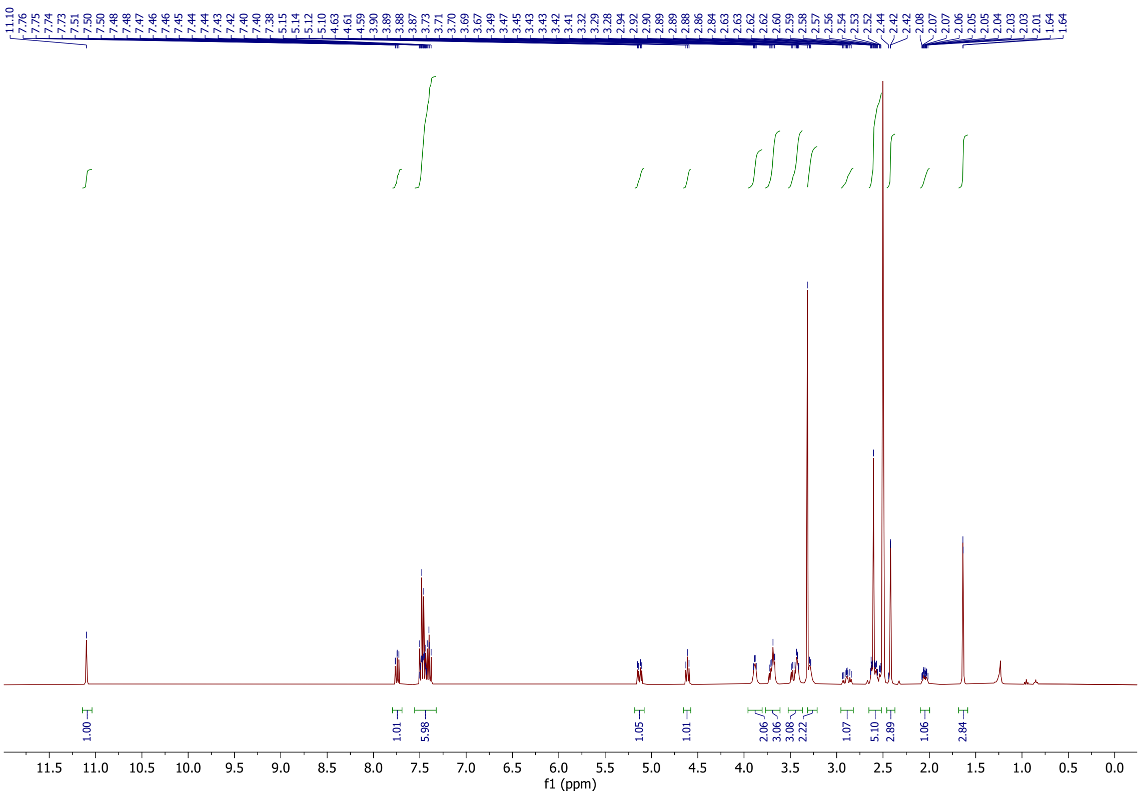

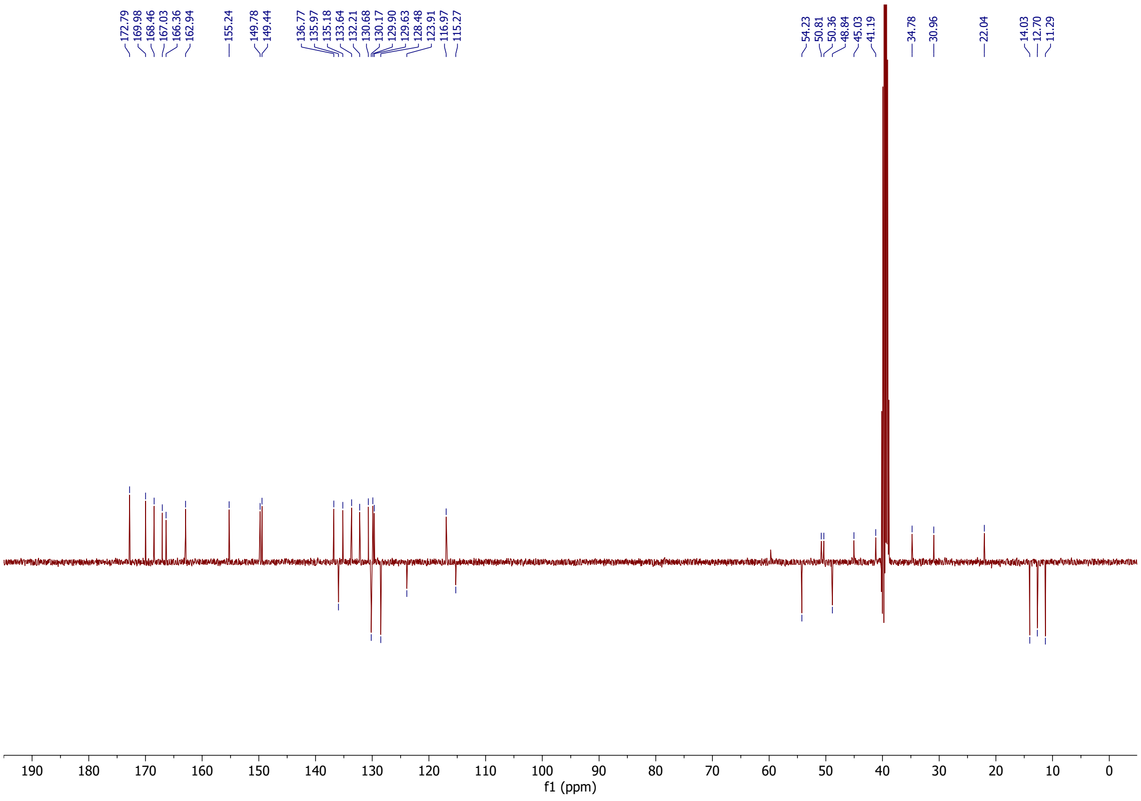

1H- and 13C-NMR spectra of compound **16**

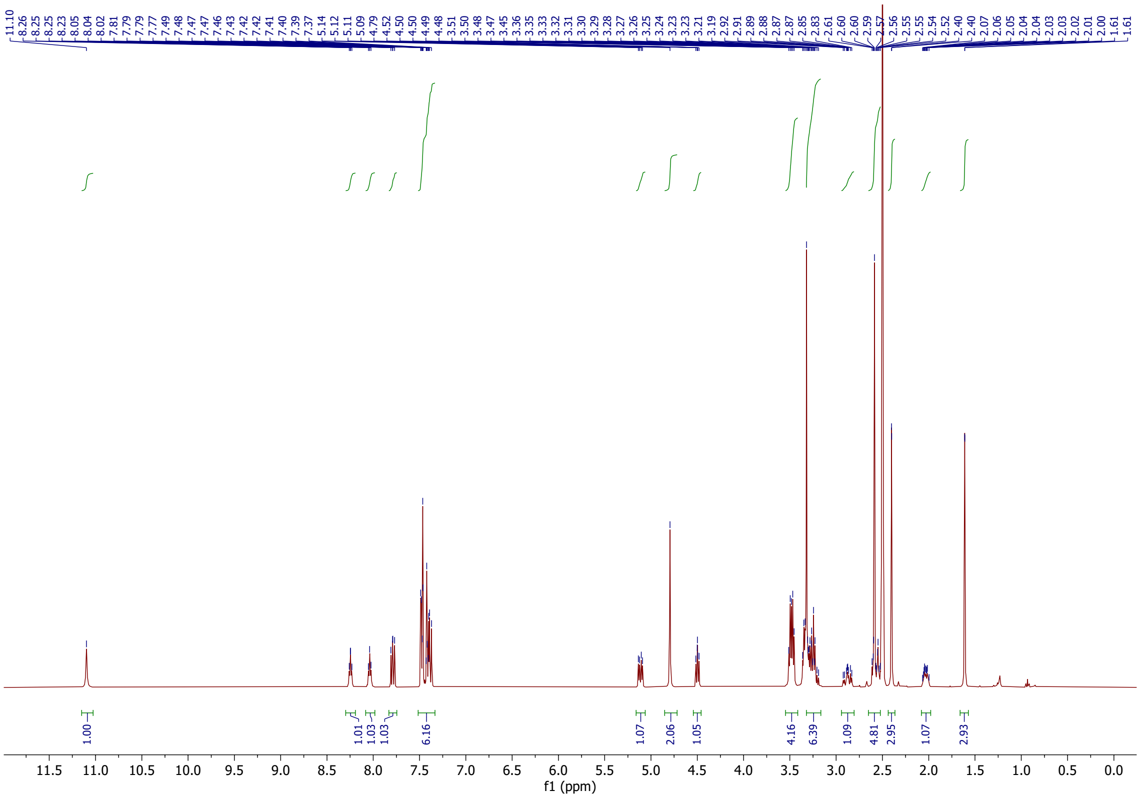

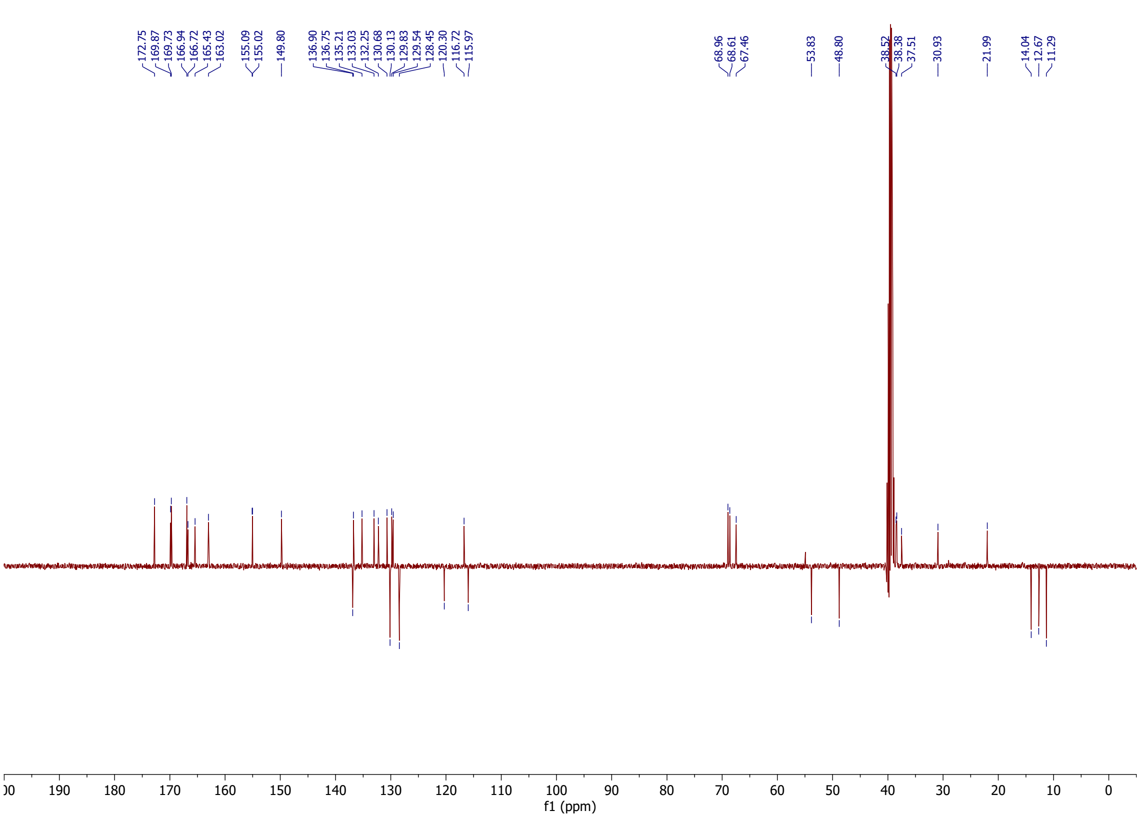

1H-NMR spectra of compound **S-9**

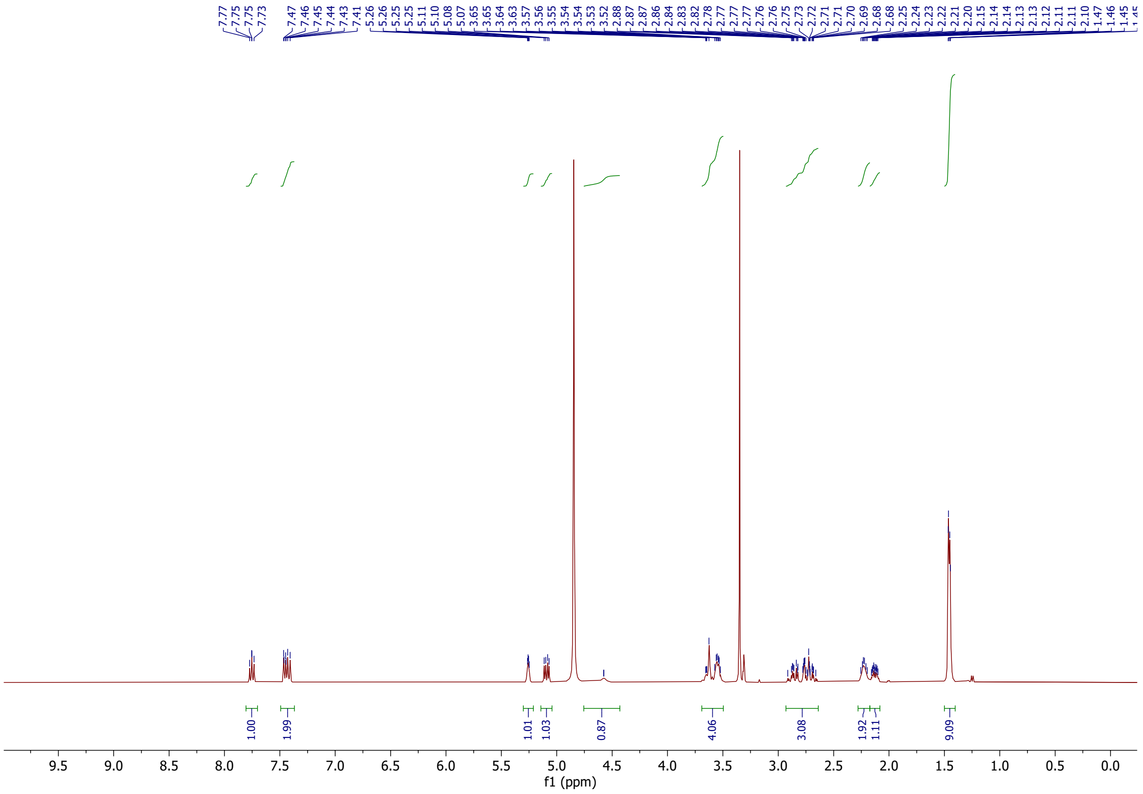

1H- and 13C-NMR spectra of compound **17**

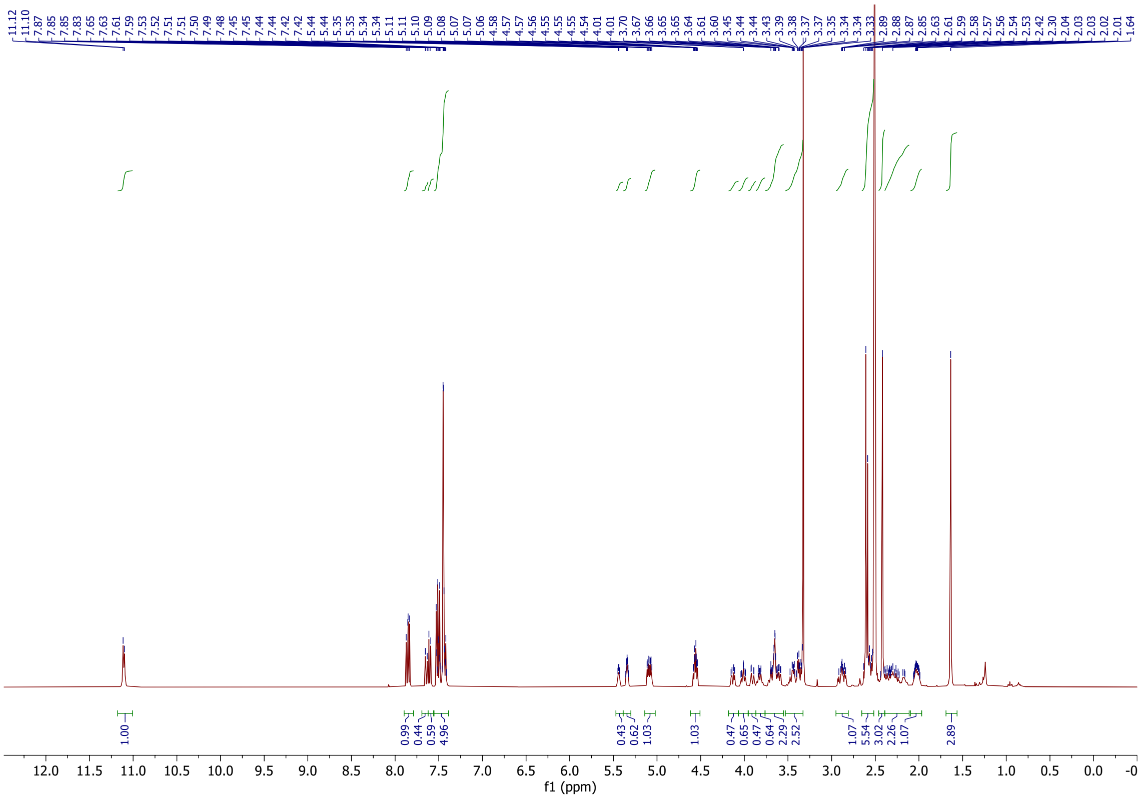

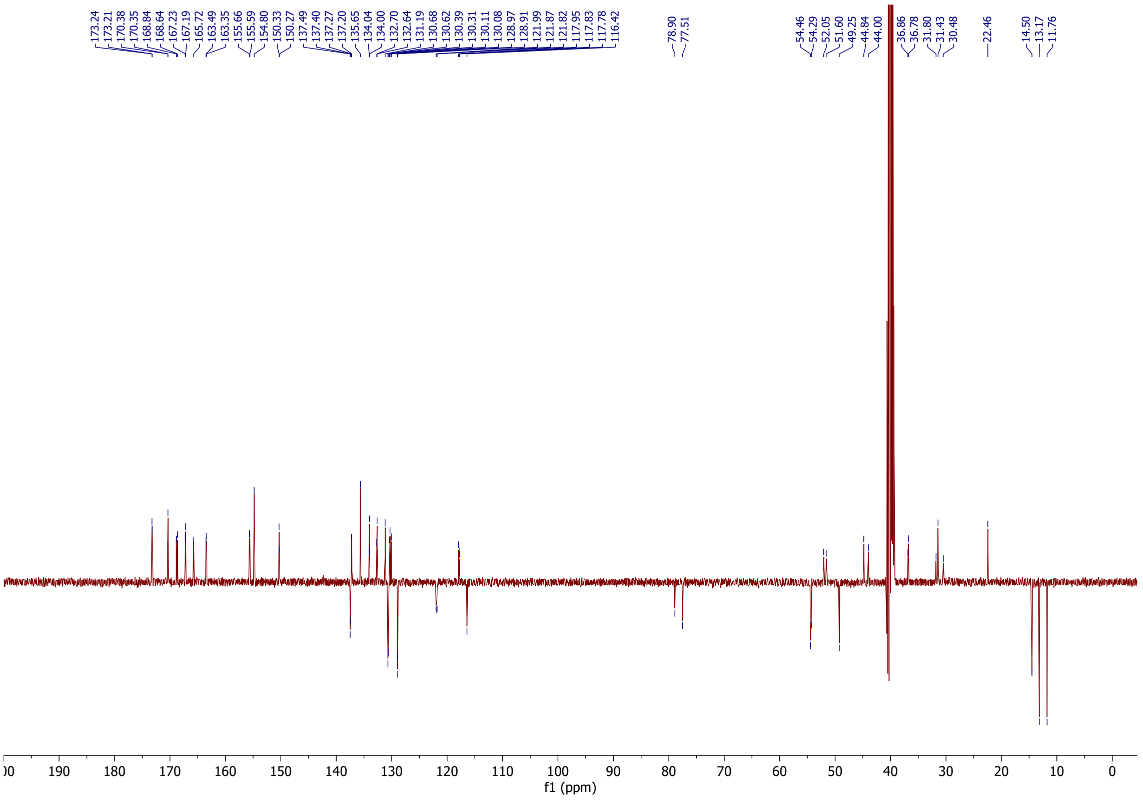
